# Proximity Interactome analyses unveil novel regulators of IRE1α canonical signaling

**DOI:** 10.1101/2024.10.27.620453

**Authors:** Simon Le Goupil, Céline Philippe, Hadrien Laprade, Mathéo Lode, Rachel Boniface, Diana Pelizzari-Raymundo, Kurt Dejgaard, Gregor Jansen, Celia Limia, Claudio Hetz, S. Jalil Mahdizadeh, Luc Negroni, Marc Aubry, Jean-Ehrland Ricci, Leif A. Eriksson, Eric Chevet

## Abstract

The unfolded protein response (UPR) is a key adaptive pathway that controls endoplasmic reticulum (ER) homeostasis. The UPR is transduced by three ER-resident sensors of ER homeostasis disruption in the lumen of this compartment. They trigger select downstream signaling pathways in the cytosol and nucleus. Among them, IRE1α (referred to as IRE1 hereafter), a type I transmembrane protein, senses accumulation of improperly folded proteins in the ER lumen and transduces signals through both kinase and endoribonuclease (RNase) activities in the cytosol. IRE1 catalyzes *XBP1* mRNA unconventional splicing and RNA degradation (Regulated IRE1 Dependent Decay, termed RIDD). Recent studies have reported that IRE1-dependent protein-protein interactions (PPi) drive additional non-canonical IRE1 functions. Herein, we define the IRE1 signalosome as a list of IRE1 binding partners (direct or not) which alter IRE1 signaling towards XBP1 mRNA splicing and RIDD. Here we determined the IRE1 *in situ* interactome using BioID, putatively connecting IRE1 to previously unrecognized cellular functions. In addition, we link the binding of several IRE1 partners to the regulation of its RNase. Furthermore, we identify HNRNPL as an IRE1-interacting partner, previously unrecognized, which stabilizes IRE1 under basal conditions by counteracting ERAD-mediated degradation. Overall, the characterization of the IRE1 signalosome not only reveals the multi-faceted control of IRE1 RNase activity and stability by its interacting partners and allow us to discuss putative additional IRE1 regulators and cellular functions based on the nature of its interactome and its localization.

## Introduction

The endoplasmic reticulum (ER) is the first compartment of the secretory pathway. In addition to its functions in regulating calcium and lipid homeostasis, the ER plays a predominant role in protein synthesis, productive folding and trafficking (Ashby and Tepikin, 2001; Fagone and Jackowski, 2009; Braakman and Hebert, 2013; Almanza et al., 2019). Indeed, approximately one-third of the proteome passes through the ER to acquire proper folding before reaching its final destination (Ghaemmaghami et al., 2003). The ER lumen exhibits a calcium-rich, oxidative environment populated by ER-resident enzymes and foldases for proper protein folding (Hanson et al., 2009). In addition, it provides a protein quality control system responsible for either retaining misfolded proteins in the ER to allow further folding cycles or promoting the degradation of terminally misfolded proteins through ER-associated degradation (ERAD) (Olzmann et al., 2013). ER homeostasis is disrupted when demand for protein processing in the ER exceeds its protein folding or degradation. This leads to an accumulation of misfolded proteins in this compartment, a situation known as ER stress (Schröder and Kaufman, 2005). To cope with ER stress, organisms/cells have evolved a signaling pathway called the Unfolded Protein Response (UPR). The UPR is triggered upon ER stress primarily to restore ER homeostasis (adaptive UPR), however, if this fails, the UPR engages cells in a death path (terminal UPR) (McGrath et al., 2021). Over the years, many studies have demonstrated the involvement of UPR signaling in a variety of diseases, such as cancer or neurodegenerative diseases (Hughes and Mallucci, 2019; Madden et al., 2019).

The UPR is transduced by three ER-resident transmembrane sensors called PKR-like Endoplasmic Reticulum Kinase (PERK), Activating Transcription Factor 6 (ATF6) and Inositol Requiring Enzyme 1 (IRE1). PERK is a type I integral membrane protein with a luminal sensing domain and a cytosolic kinase domain. Activated PERK phosphorylates eukaryotic initiation factor 2 alpha (eIF2α) at serine 51, thereby decreasing overall translation and therefore ER protein load (Hetz et al., 2020). Phosphorylated eIF2α also allows translation of selected mRNAs exhibiting µORF (upstream Open Reading Frame) such as that encoding the transcription factor ATF4 (Vattem and Wek, 2004). Major targets of ATF4 include genes involved in redox homeostasis, amino acid metabolism and autophagy (B’chir et al., 2013; Harding et al., 2003) as well as its own downregulation pathway (Novoa et al., 2001). Upon ER stress, ATF6, a type II ER transmembrane protein, is transported to the Golgi apparatus by a COPII-dependent mechanism, where it is cleaved by the site 1/2 proteases (S1P and S2P) leading to the release of the ATF6 cytosolic domain (ATF6f) (Haze et al., 1999). ATF6f primarily induces the expression of genes encoding for of ER chaperones/foldases, leading to an increase in productive protein folding in the ER (Wu et al., 2007; Shoulders et al., 2013).

The most conserved UPR transducer is IRE1α, discovered in yeast and mammals in the 1990’s (Mori et al., 1993; Welihinda et al., 1998). IRE1α is an ER-resident type I transmembrane protein that contains both BiP-binding and misfolded protein-binding domains in the ER lumen as well as kinase and endoribonuclease (RNase) catalytic domains in the cytosol. Upon ER stress, IRE1α (hereafter referred to as IRE1) oligomerizes and trans-autophosphorylates to trigger RNA cleavage that is associated with two distinct outcomes managed by the RNase domain (Cox et al., 1993; Mori et al., 1993). First, X-Box Binding Protein 1 (*XBP1*) mRNA undergoes an unconventional splicing, in which a 26 nucleotide intron is removed from *XBP1* mRNA through the conjugated action of IRE1 and of a tRNA ligase, called RtcB, to form the spliced XBP1 (*XBP1s*) mRNA (Jurkin et al., 2014). *XBP1s* mRNA is translated into a transcription factor that induces the expression of genes involved in lipid biogenesis, ER chaperones, and ERAD (ER Associated Degradation) (Sidrauski and Walter, 1997; Lee et al., 2003, 2008; Yamamoto et al., 2008). Second, Regulated IRE1 Dependent Decay (RIDD) of RNA, which leads to the degradation of selected RNA (mRNA, rRNA, miRNA) (Hollien and Weissman, 2006; Upton et al., 2012). In addition to these catalytic activities, other IRE1 functions rely on protein-protein interactions (PPi) (Le Goupil et al., 2024). The concept of the UPRosome was initially proposed where IRE1 is envision as a scaffold where different proteins assemble to regulate its activity and mediate the crosstalk with other signaling pathways and cellular processes (Hetz and Glimcher, 2009).

Herein, we performed a comprehensive analysis of the IRE1 interactome through BioID and computational analysis that highlights the expected but also unexpected partners of IRE1, defining the IRE1 proximal signalosome as the proteins that interact with IRE1 *in situ* and modulate its catalytic activity IRE1 signalosome will be seminal for future work on IRE1 regulations and functions. As proof of concept, we further describe a new interactor of IRE1, HNRNPL (Heterogeneous nuclear ribonucleoprotein L) that regulates IRE1 protein expression through protein stabilization. We also discuss new functional interactions between IRE1 signaling and specific molecular machines associated with RNA processing.

## Results

### The reference IRE1 signalosome includes modulators of ER homeostasis

Human IRE1 is composed of an N-terminal luminal sensing domain, a short single-spanning transmembrane domain, and a cytosolic domain consisting of two main parts: a kinase domain proximal to the membrane (AA 571-832) and a C-terminal RNase domain (AA 835-963) (**Fig. 1A**). Both luminal and cytosolic domains have been shown to interact with other proteins (Le Goupil et al., 2024), some of which appear to regulate IRE1 signaling properties and functions. To explore the known PPi-based functions of IRE1 and to predict additional functions of IRE1, we first aimed to define its “reference” interactome. To do this, we performed a meta-analysis with data from public PPi repositories (Biogrid, IntAct, String, APID) and published dataset (Acosta-Alvear et al., 2018) (see Materials and Methods). Noteworthy that most of the interactors found in this analysis were identified using either targeted co-immunoprecipitation or IRE1 immunoprecipitation followed by mass spectrometry (MS) sequencing. The reference IRE1 protein interaction network resulting from this meta-analysis contained 567 proteins (**Fig. 1B**), and KEGG enrichment analysis revealed the terms “Protein processing in the ER” and “Apoptosis” as major clusters in the network, both of which are known IRE1-associated functions. In addition, we found enriched terms in the network that linked IRE1 to ribosomal proteins and ribosome biogenesis, as well as processes associated with genome maintenance (**Fig. 1B, C**). Only a limited number of studies described links between IRE1 and the translation machinery (Acosta-Alvear et al., 2018; Belyy et al., 2022; Gómez-Puerta et al., 2022) or DNA damage and repair (Dufey et al., 2020), and therefore these remain largely unexplored, suggesting uncharacterized functions for IRE1. We next aimed to identify partners of IRE1 that are also regulators of its signaling. Therefore, we intersected the IRE1 reference interactome with hits that alter XBP1 mRNA splicing, as previously identified in a CRISPR screen (Adamson et al., 2016) and two siRNA screens (Papaioannou et al., 2022; Yang et al., 2018) aiming at identifying hits that alter *XBP1* mRNA splicing, a read-out of IRE1 activity. These data were intersected with the IRE1 reference interactome to identify regulators of IRE1 signaling that are also in complex with IRE1 (**Fig. 1D**). As a result, 77 proteins were found to be shared between the reference interactome and the CRISPR/siRNA screens. We defined this list of 77 proteins as the “IRE1 reference signalosome”, and assembled them into a PPi network (**Fig. 1E**). The most enriched Gene Ontology (GO) terms of this reference signalosome highlighted processes related to ERAD, UPR, protein folding, and trafficking, all of which are indicative of canonical IRE1 functions and related to ER homeostasis (**Fig. 1F**). This indicates that ER homeostasis may be regulated by means involving IRE1-PPi network through the regulation of IRE1 activity. However, since this reference interactome mainly rely on co-immunoprecipitation and immunoprecipitation coupled with MS, one main limitation of these approaches resides in the loss of cell compartmentalization which can generate interactions that are not physiologically relevant.

**Figure 1:**
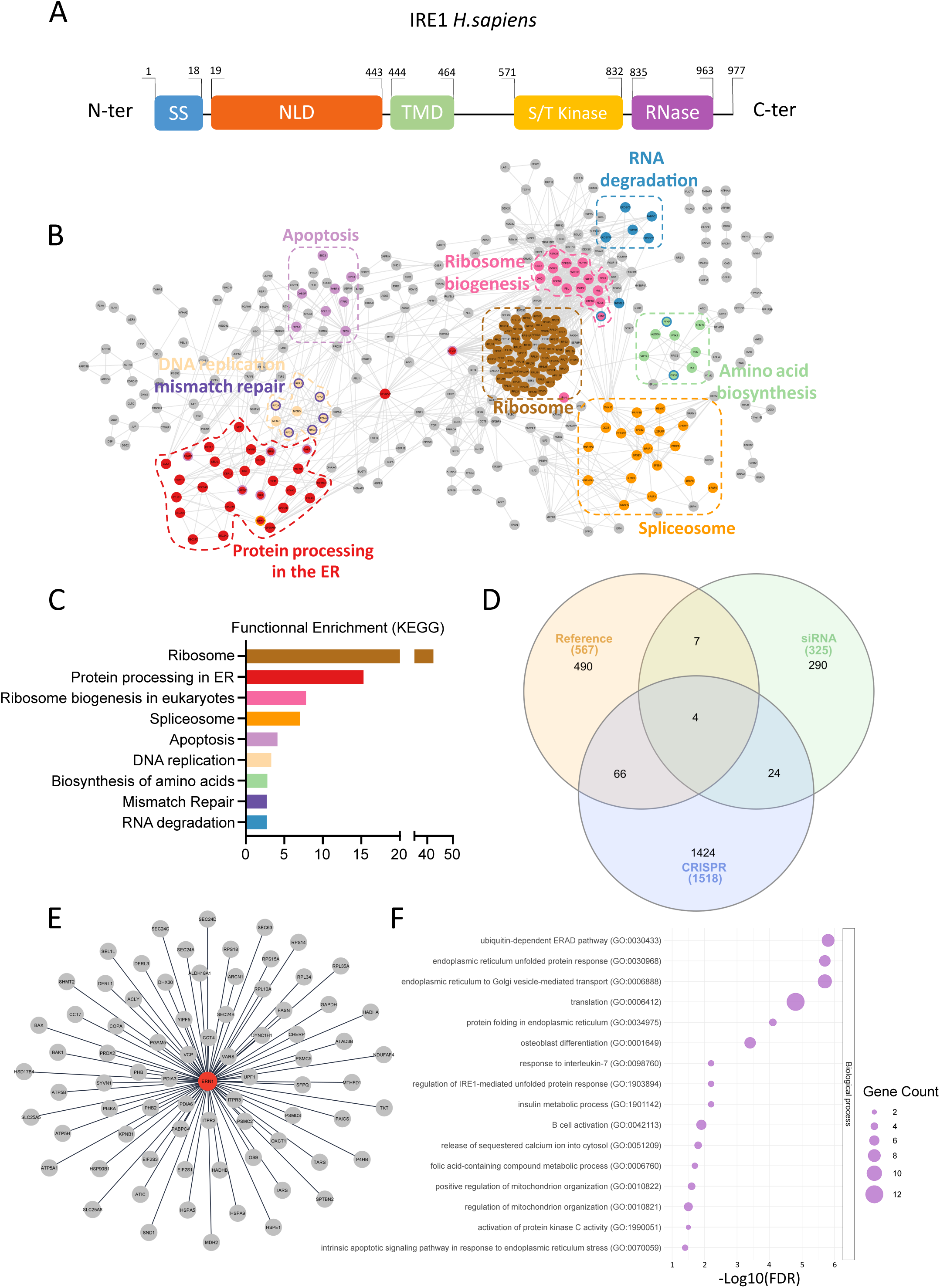
IRE1 Reference interactome and associated signalosome. **A)** Schematic representation of IRE1 sequence. **B)** PPi network made with Cytoscape depicting the 567 proteins of IRE1 reference interactome, with colored clusters defined by KEGG in (C). **C)** KEGG functional enrichment of the IRE1 reference interactome. **D)** Venn diagram crossing the lists of IRE1 reference interactome, siRNA and CRISPR screenings, defining the IRE1 reference signalosome. **E)** PPi network made with Cytoscape depicting the 77 proteins from the IRE1 reference signalosome. **F)** Gene ontology enrichment of the IRE1 reference signalosome.

### Validation of the IRE1 BioID construct

Most of the data used to generate this IRE1 reference interactome were collected using either targeted co-immunoprecipitation or IRE1 immunoprecipitation followed by mass spectrometry (MS) sequencing. The main limitation of these approaches resides in the loss of cell compartmentalization which can generate interactions that are not physiologically relevant. Furthermore, weak/transiently interacting partners may not be detected through these approaches. Therefore, we aimed to establish an IRE1 interactome of cytosolic components using a method that directly screens PPi in living cells. We chose a proximity-dependent labelling approach, the BioID, which allows for the maintenance of cell compartmentalization (Sears et al., 2019). The biotin ligase BirA* gene was fused to the C-terminal end of the *IRE1* gene along with a Flag tag forming the *IRE1-BirA\** fusion gene (**Fig. 2A**). To validate the correct localization and activity of this fusion protein, HeLa cells were transiently transfected with *IRE1-BirA\** and treated with or without biotin prior to immunofluorescence analysis. As expected, the Flag signal colocalized with the ER membrane marker calnexin (CNX) as shown by the merge, indicating that IRE1-BirA* is correctly localized at the ER membrane (**Fig. 2B**). Biotinylated proteins stained using FITC-conjugated streptavidin, were detected exclusively in cells treated with biotin, indicating that exogenous biotin is necessary for IRE1-BirA*-dependent biotinylation. In addition, biotinylated proteins mainly colocalized calnexin indicating that most of IRE1-BirA* dependent biotinylation occurred at the ER and nuclear periphery (**Fig. 2B)**.

**Figure 2:**
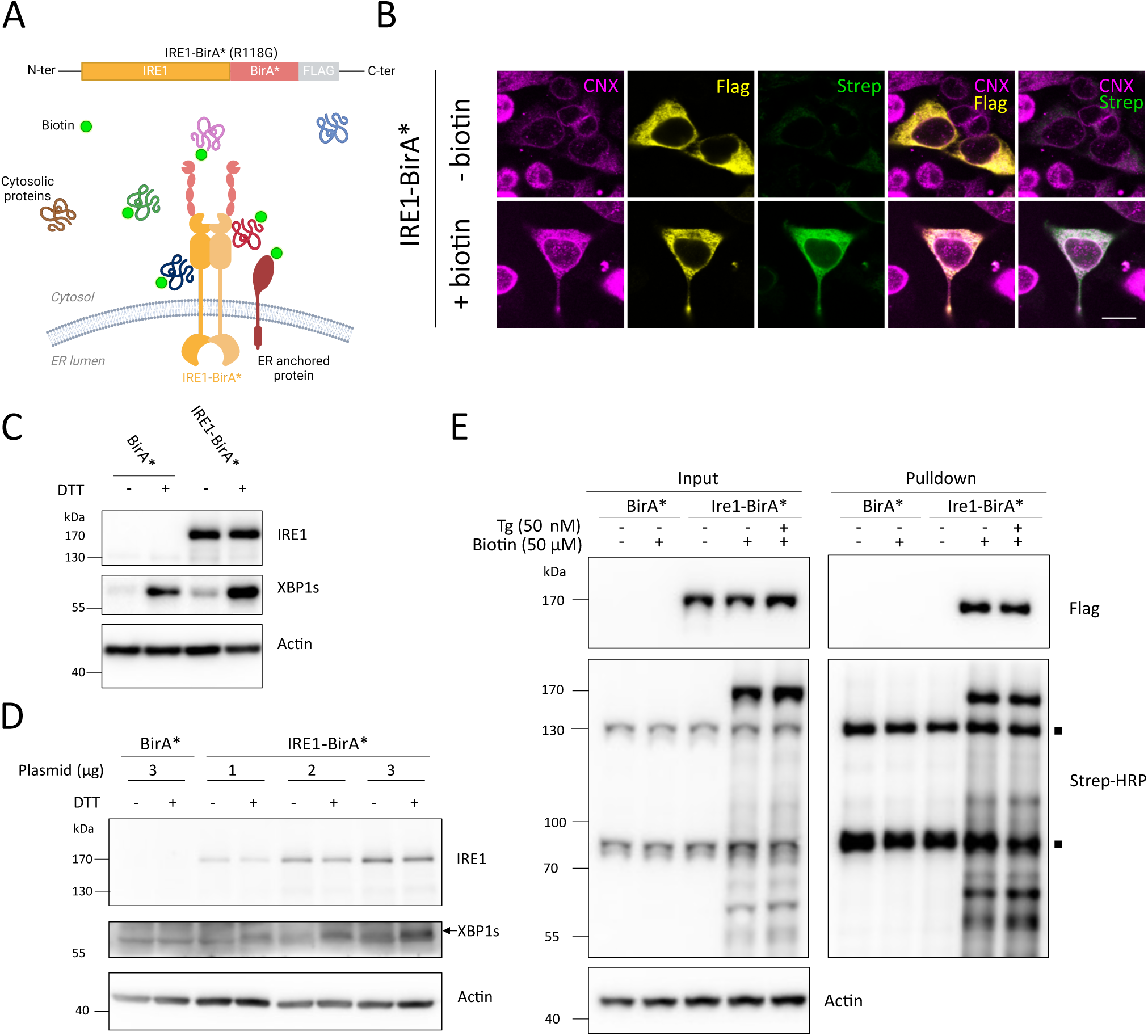
Validation of the IRE1-BioID construct. **A)** Schematic representation of the IRE1-BirA* sequence (top) and schematic representation of IRE1-BirA* dependent biotinylation (bottom). The green dots represent biotin residues, added on IRE1-interacting protein in a 10 nm radius. **B)** Immunofluorescence images of HeLa cells after 48 hours IRE1-BirA* transfection and treatment with or without biotin (50 µM) for 24 hours. Cells are stained for calnexin (CNX), Flag and biotinylated proteins (Streptavidin, Strep). Scale bar = 100 µm. **C, D)** Western-Blot of IRE1-BirA* expression in transiently transfected (48 hours) HEK293T (C) or IRE1 KO A375-MA2 (D) cells treated with DTT for 4 hours. The arrow targets XBP1 specific band. One representative experiment out of three is shown. **E)** Western-Blot of IRE1-BirA* dependent biotinylation in HEK293T cells. Cells were transfected with IRE1-BirA* for 48 hours. At 24 hours, 50 µM biotin was added to the medium and 50 nM thapsigargin further added for 6 hours. Biotinylated proteins were pulled down by neutravidin magnetic beads and analyzed using HRP-coupled streptavidin. ▪ corresponds to endogenous biotinylated proteins. One representative experiment out of three is shown.

Next, we evaluated the RNase activity of IRE1-BirA*. HEK293T cells transiently expressing IRE1-BirA* exhibited higher basal XBP1s than control cells. In addiation, XBP1s was further increased upon ER stress induction by dithiothreitol (DTT) (**Fig. 2C**). Hence, IRE1-BirA* RNase is active and sensitive to ER stress (**Fig. 2C**). To exclude a potential role of endogenous IRE1, we reproduced this experiment in the metastatic melanoma cell line A375-MA2 knocked out (KO) for *IRE1* (**Fig. S1**). A375-MA2 *IRE1* KO cells were rescued by increasing amount of IRE1-BirA*. Indeed, the expression of IRE1-BirA* which correlated with a progressive and DTT-sensitive increase in XBP1s induction (**Fig. 2D**). These data confirmed that IRE1-BirA* is functional for *XBP1* mRNA splicing. Selective IRE1-BirA*-mediated protein biotinylation was also observed by western-blot, in addition to endogenous carboxylase biotinylation (Tong, 2013) (**Fig. 2E**), consistent with the result obtained in **Fig. 2B**. Biotinylated proteins were enriched using streptavidin-coated beads and analyzed by blotting with streptavidin-HRP. Of note, mild ER stress induced by thapsigargin did not alter the biotinylation pattern observed in IRE1-BirA* transfected cells (**Fig. 2E**). Taken together, these results demonstrate that IRE1-BirA* maintains basal and ER-stress inducible XBP1 splicing activity and is able to biotinylate proximal interactors,

### The IRE1 *in situ* interactome

To define the IRE1 *in situ* interactome in a manner that preserves cellular compartments, samples from biotin-treated BirA* and IRE1-BirA* expressing HEK293T cells treated or not with the ER stressors tunicamycin (Tm) or thapsigargin (Tg) for 16 h were analyzed by mass spectrometry. This revealed 207 IRE1 interactors (including IRE1 itself) (**Table 1**). Most of the interactors (146) were identified exclusively under basal conditions, while 42 interactors were identified specifically under ER stress and 18 in both conditions (**Fig. 3A**), with only 13 interactors depending on the type of stress (Tg or Tm) (**Fig. S2A**). These IRE1 interactors were manually annotated according to their main cellular function. Analysis of the relative composition of each cluster revealed that under basal conditions, IRE1 interacts primarily with proteins involved in RNA processing, cytoskeleton, and metabolism (**Fig. 3B**). The interactome obtained with ER-stressed cells showed a higher proportion of interactions with chaperones and proteins involved in maintaining ER homeostasis, consistent with the canonical role of IRE1 in resolving ER stress (**Fig. 3B & S2B**). We further confirmed our observations in a systematic approach by performing GO analysis of the interactors (**Fig. 3C**).

**Table 1:**
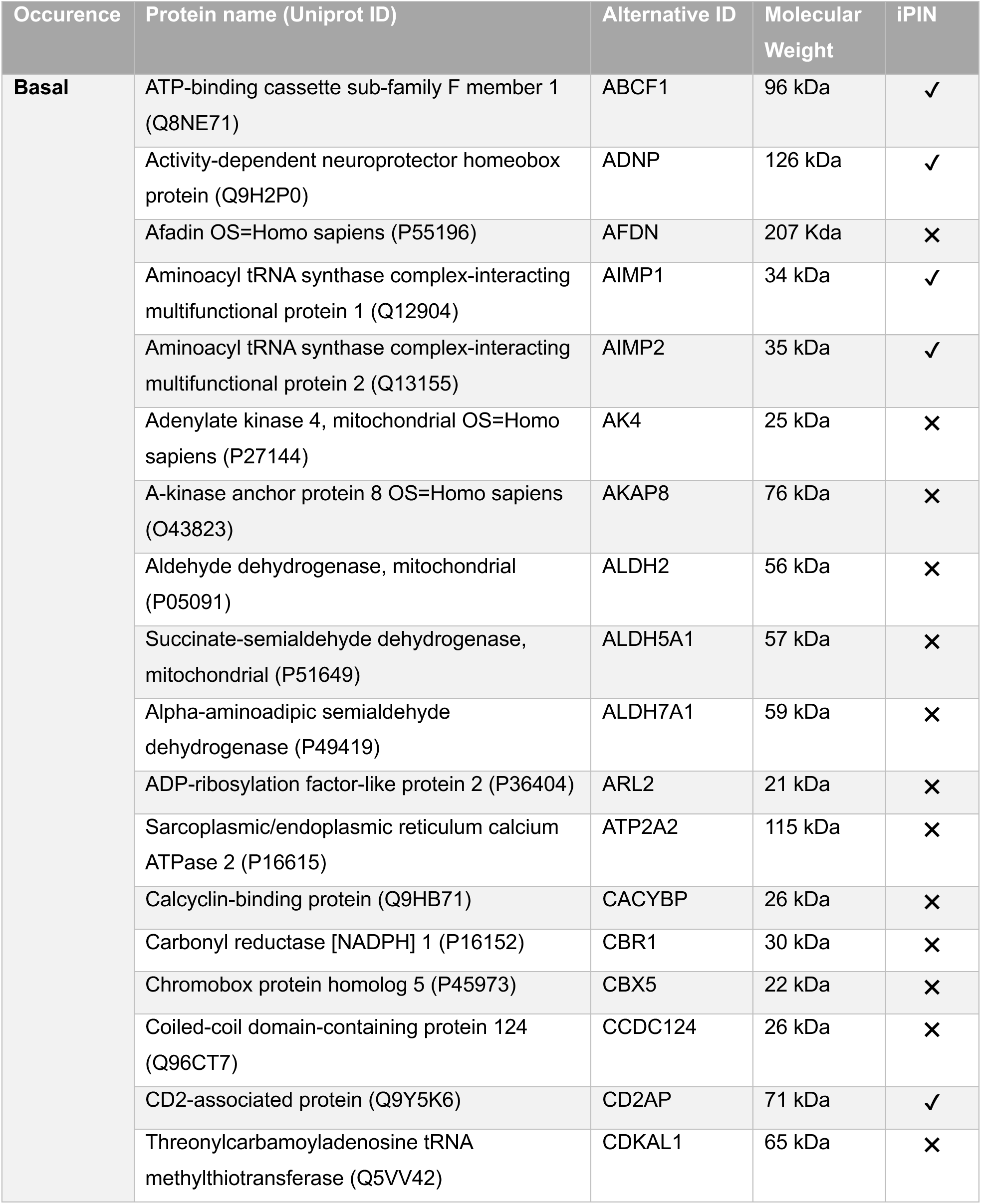
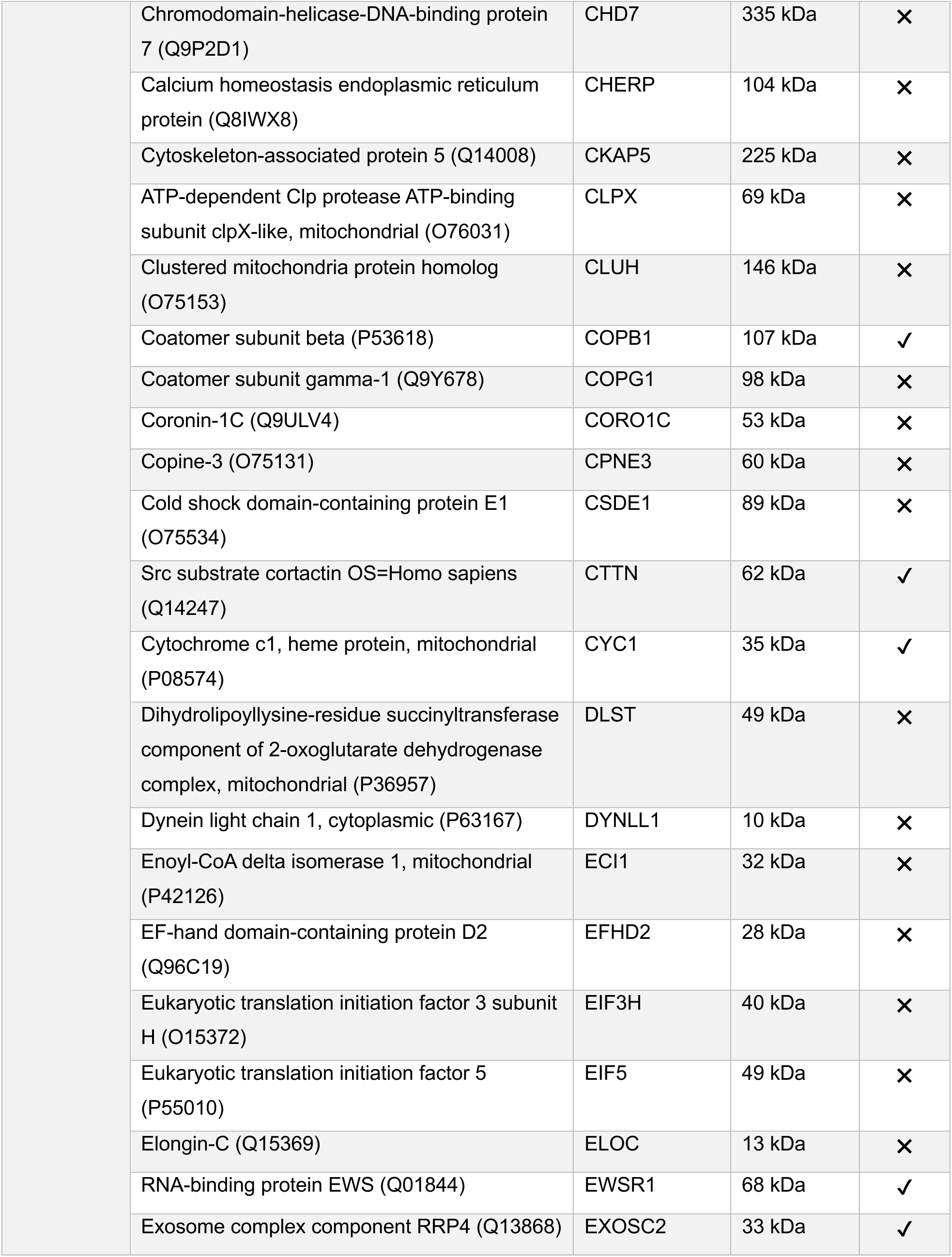
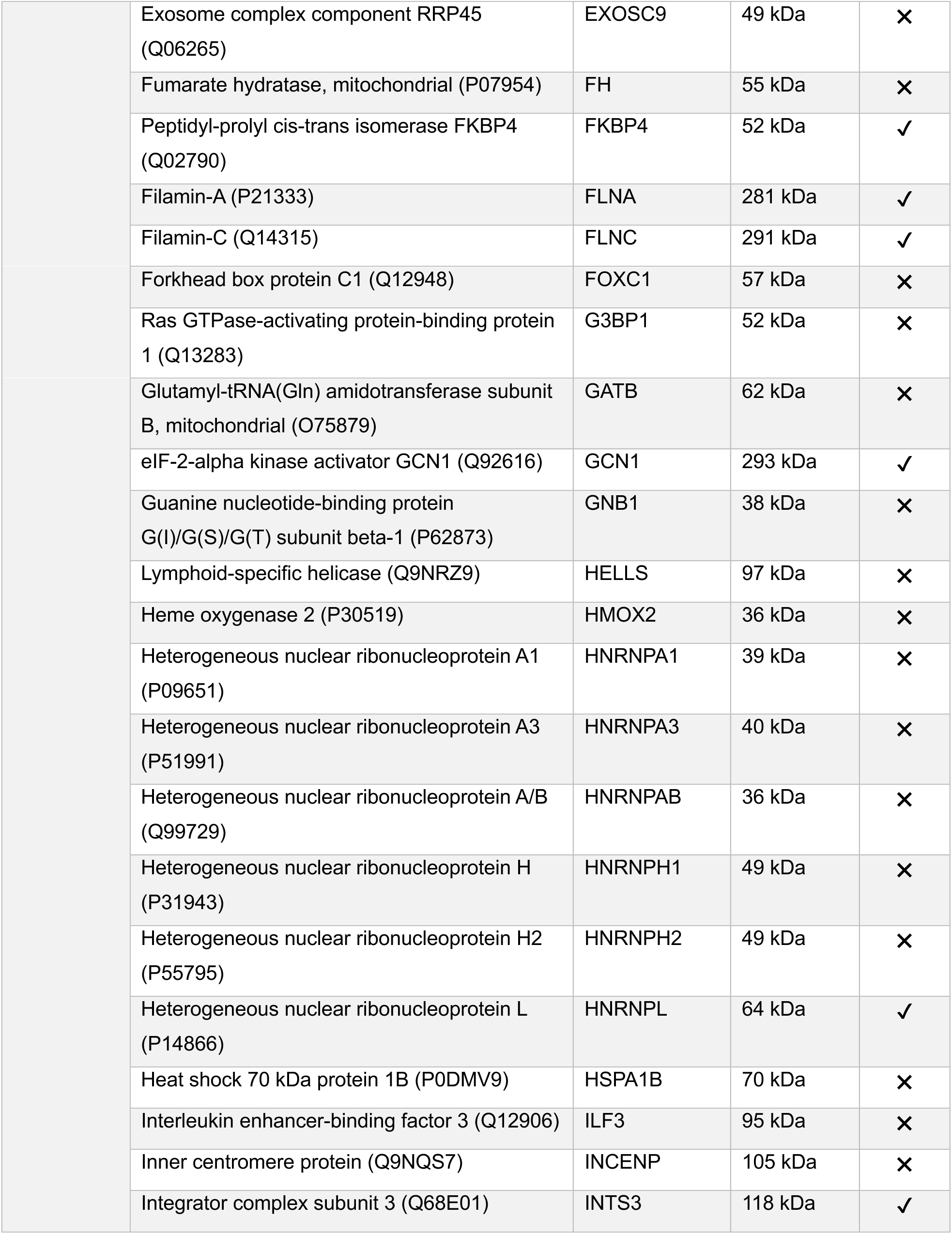
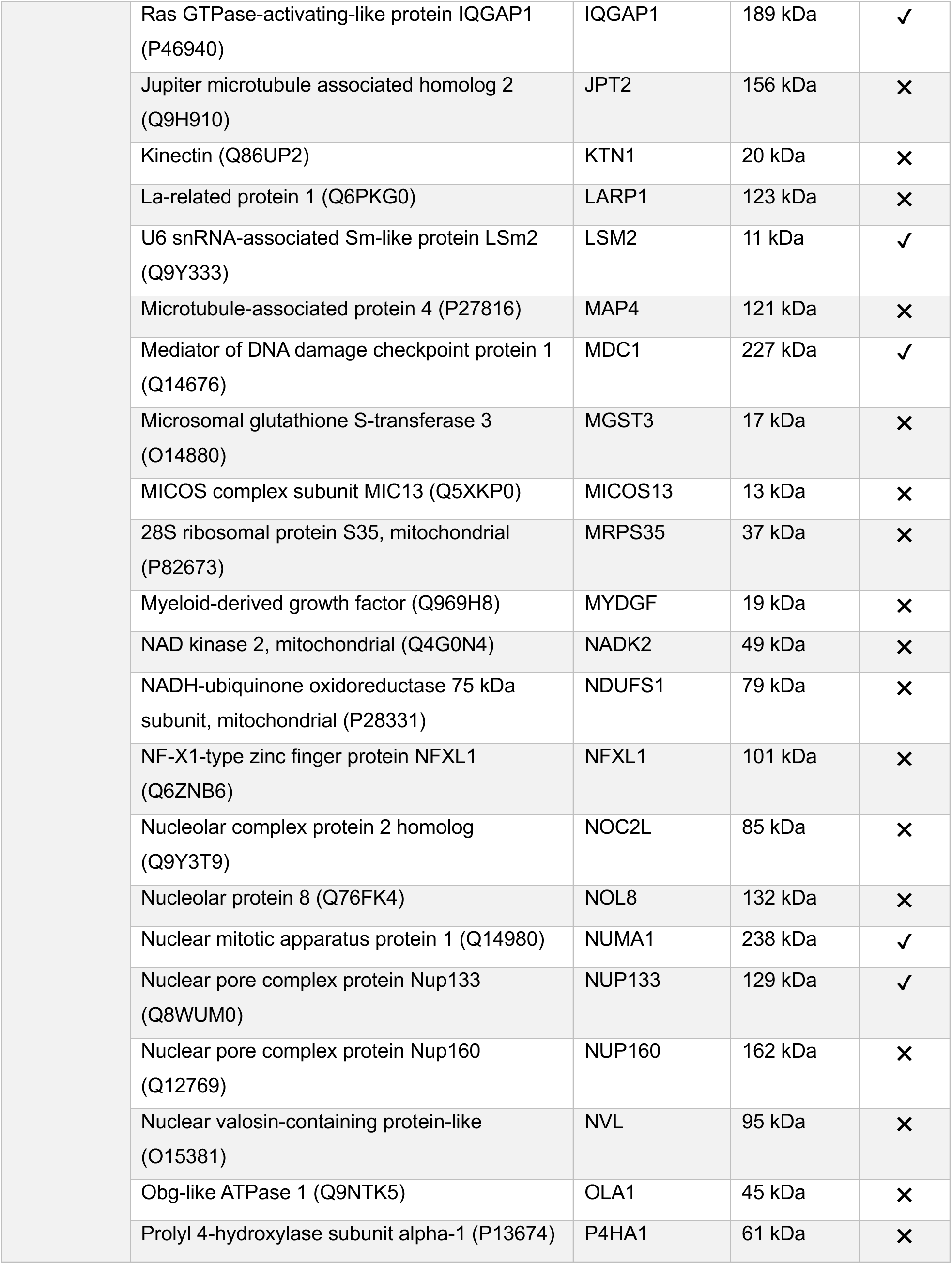
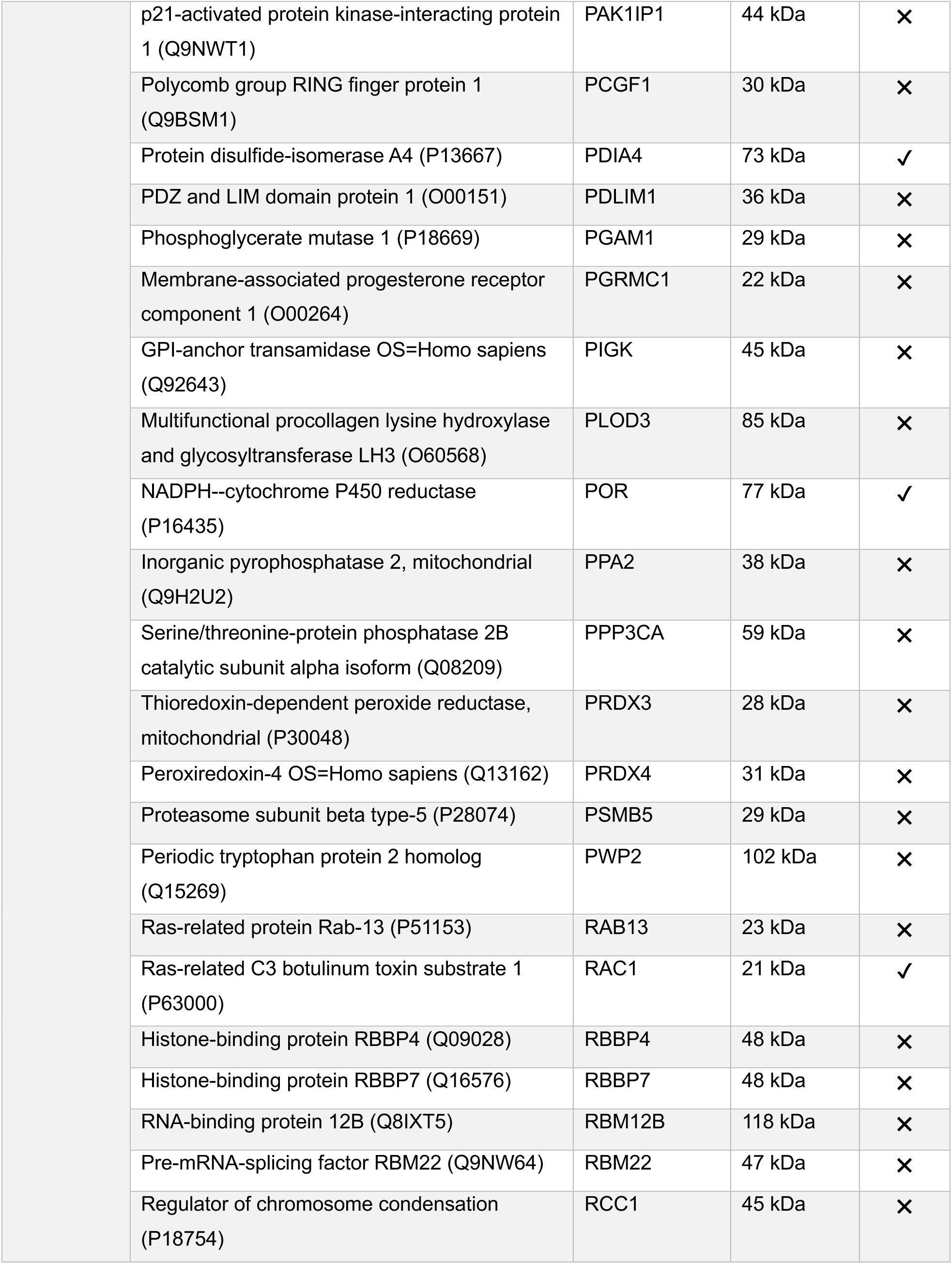
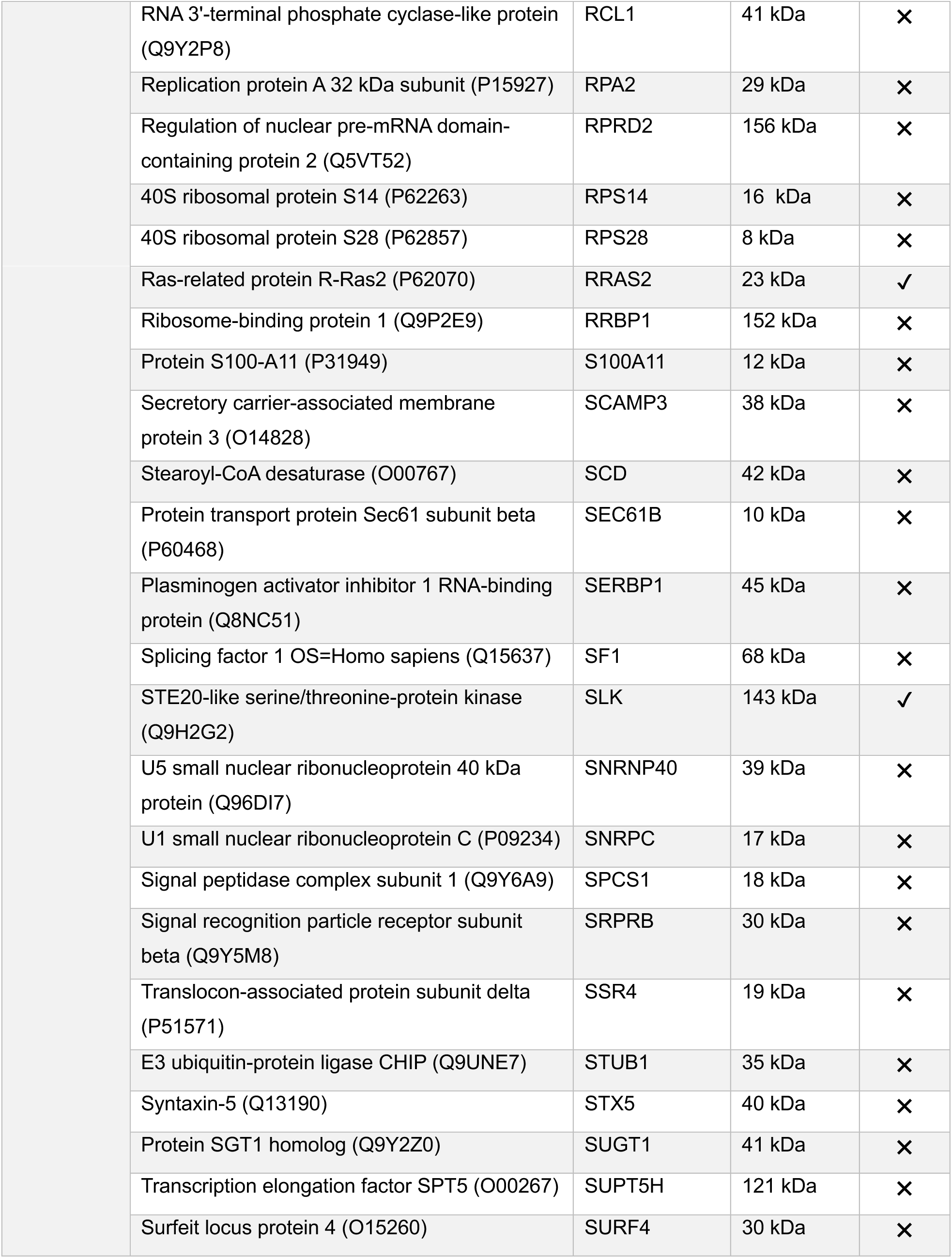
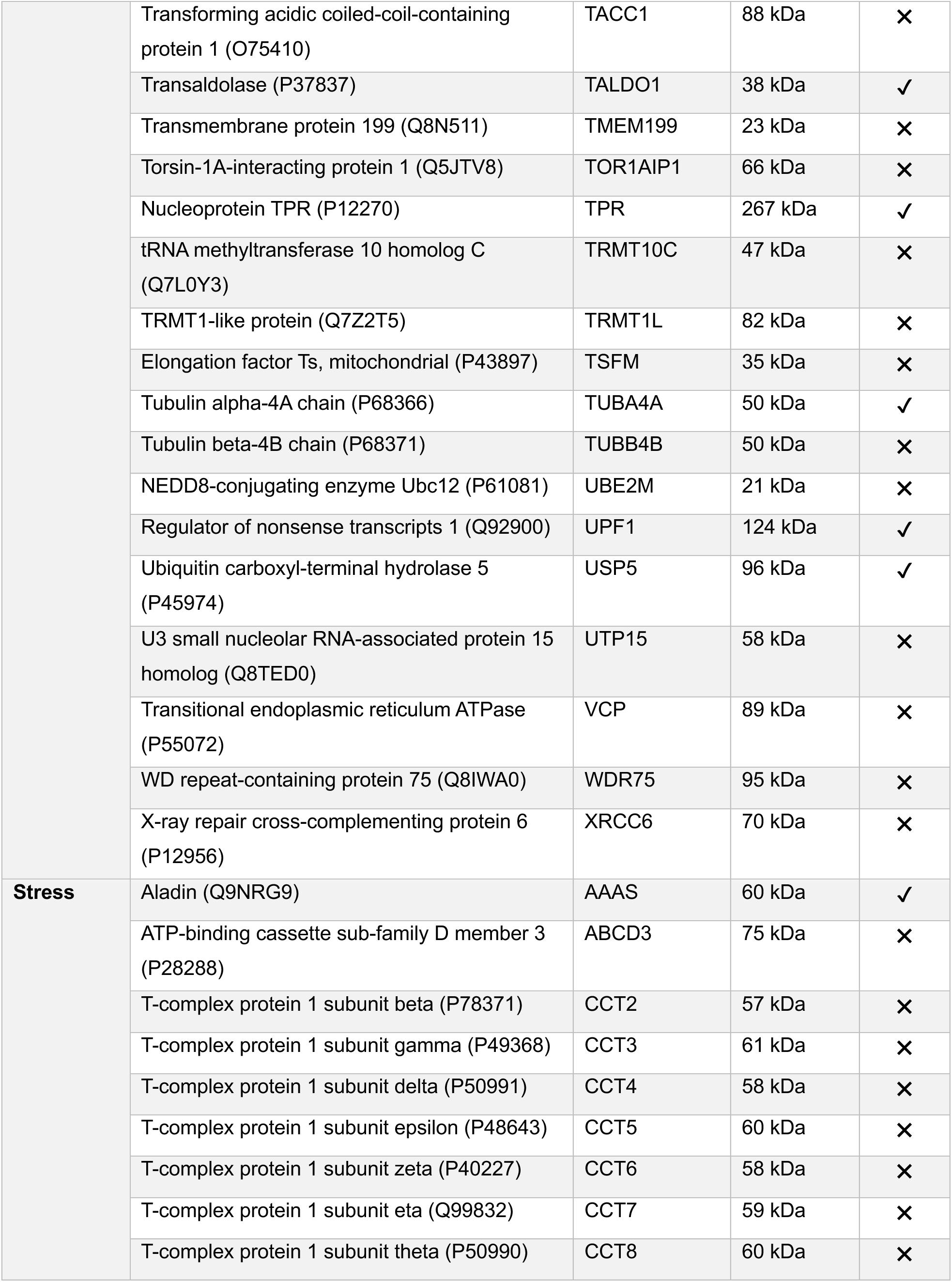
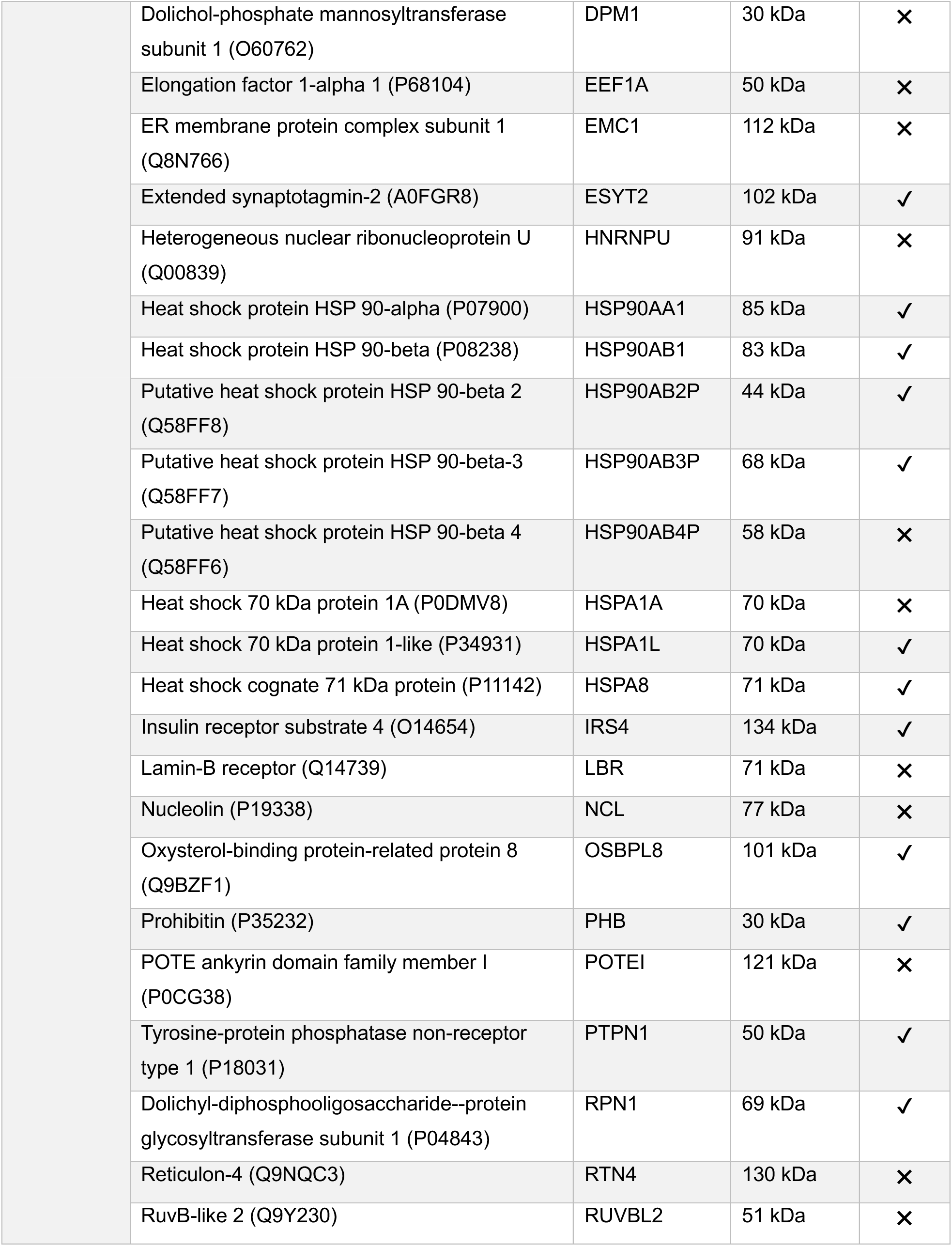
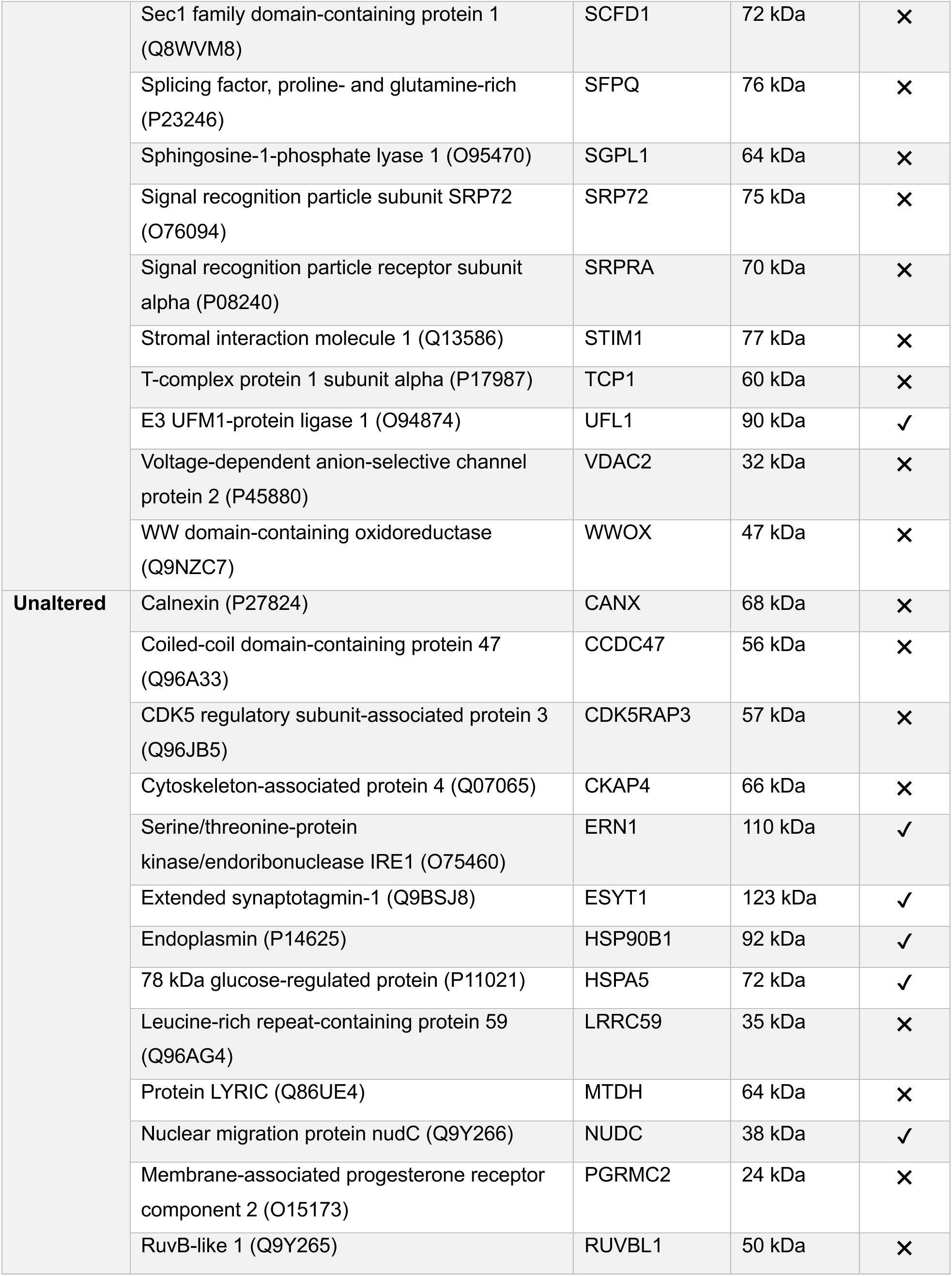
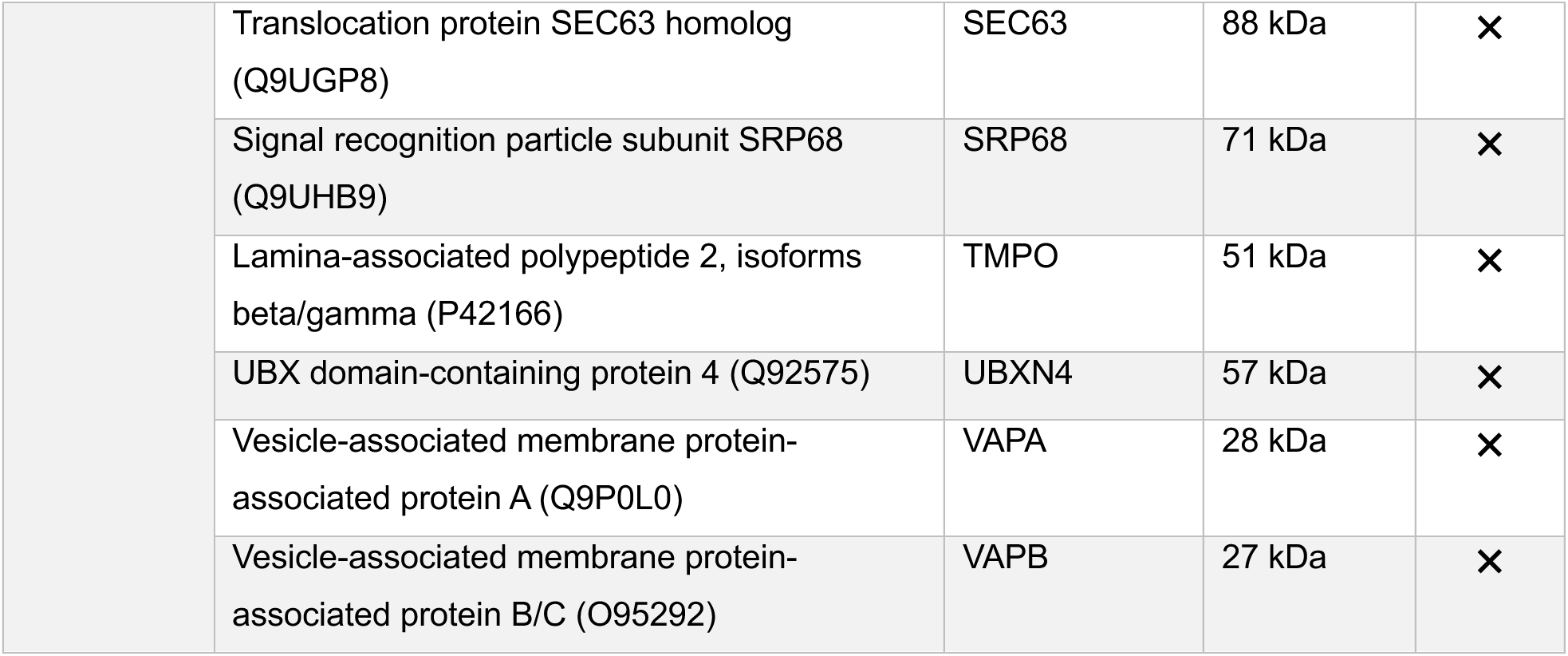
List of identified proteins resulting from the BioID interactome screen using IRE1-BirA*. Proteins listed here were only detected in IRE1-BirA* cells but not in BirA* control cells.

**Figure 3:**
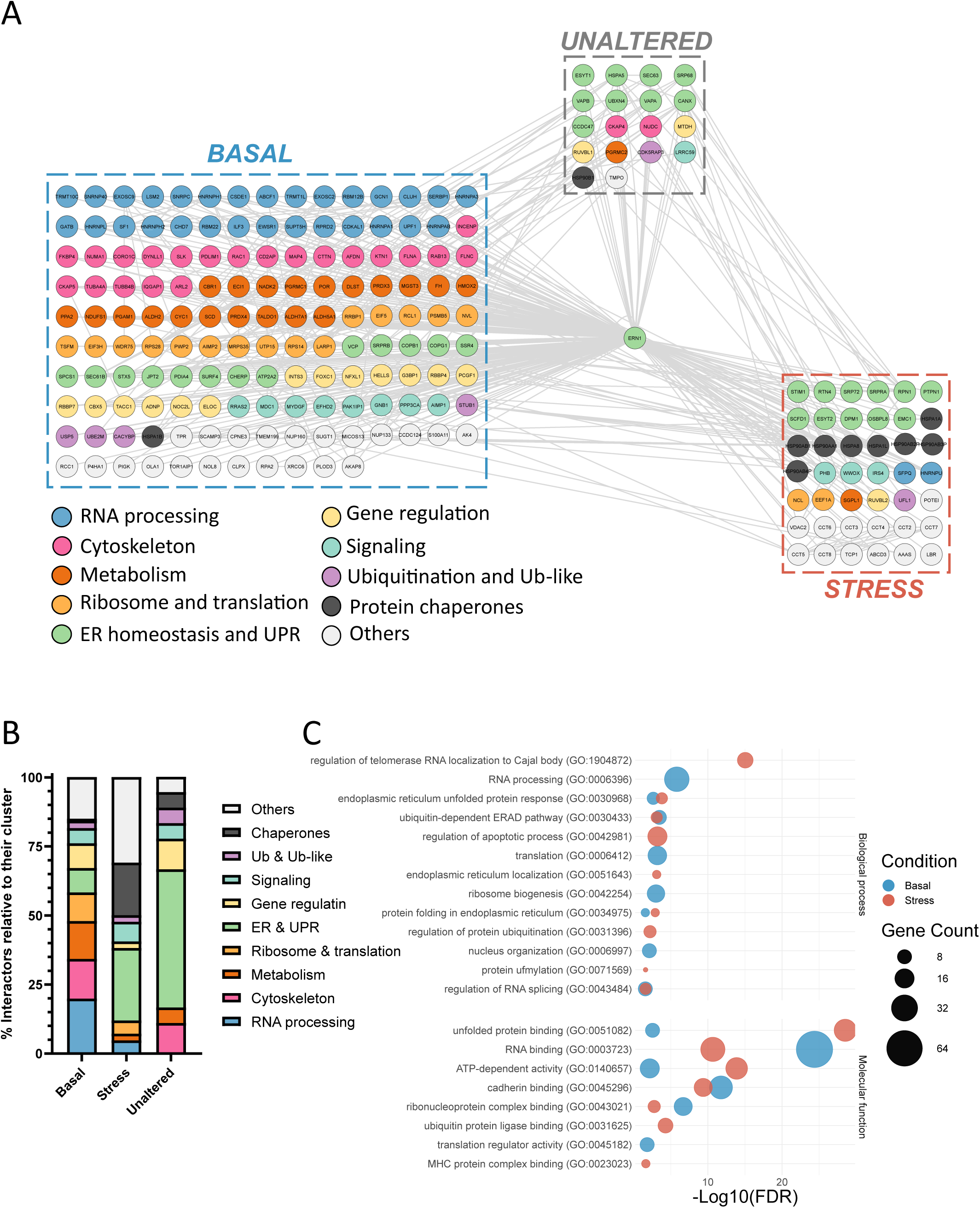
The BioID-based IRE1 interactome. **A)** PPi network of the IRE1 BioID interactome. HEK293T cells transfected with IRE1-BirA* or BirA* were treated with biotin. Cells were further left untreated or treated with ER stressors Tunicamycin (5 µg/mL) or Thapsigargin (500 nM) for 6 hours. Biotinylated proteins were isolated by streptavidin pull-down and identified by mass spectrometry. IRE1 interactome was evaluated by selecting the IRE1-BirA* specific biotinylation. Each of the 207 interactors was manually annotated according to its main function and colored accordingly. **B)** Bar graph representing the proportion of each manually curated functions within the basal, unaltered and stress clusters. **C)** Comparison of Biological process (Top) and Molecular functions (bottom) Gene Ontologies of basal and stress IRE1 BioID interactomes.

Comparing the basal and stressed conditions, we found that the basal interactome displayed higher enrichment in terms related to RNA metabolism (e.g., “RNA processing,” “translation,” “ribosome biogenesis,” “RNA binding”), whereas the stressed interactome was mainly enriched in processes associated with ER homeostasis (e.g. “ER UPR,” “regulation of apoptotic processes,” “protein folding,” “unfolded protein binding”), consistent with what was found with the manual annotation. Interestingly, when comparing the subcellular compartments of the reference (**Fig. 1B**) with the BioID IRE1 interactome, we found that the latter showed a greater compartmental diversity (**Fig. S2C**), as expected, provided that the ER forms contact with a large diversity of other compartments. Remarkably, the intersection between the IRE1 reference interactome (**Fig. 1B**) and the IRE1 BioID interactome indicated 49 shared proteins (**Fig. S2D, left**), mostly related to the UPR and protein folding (e.g., “regulation of protein stability”, “chaperone-mediated folding”, response to unfolded protein”) (**Fig. S1D, right**).

Next, we wondered whether IRE1 may share common interactors with the UPR sensor PERK, as already shown for example for their shared association with FLNA and the cytoskeleton (Urra et al., 2018; van Vliet et al., 2017). Thus, we compared our results with an available BioID interactome for PERK (Sassano et al., 2021). This revealed 43 common proteins (**Fig S2E, left**), mostly associated with functions related to ER homeostasis (e.g., “response to ER stress”, “protein folding”, “ERAD”) (**Fig S2E, right**). In addition, we compared the IRE1 BioID interactome with a previously published IRE1β interactome (Cloots et al., 2023). IRE1β shares the same features as IRE1α but differs in its RNase kinetics and tissue expression, which is restricted to mucosal epithelium for IRE1 β. We found 10 common interactors between the two homologues, mostly chaperones (e.g., HSP90B1, HSPA5, HSPA8 PDIA4) (**Fig. S2F**). Interestingly, biological processes such as cytoskeleton, RNA metabolism, translation were not enriched in the common interactome, suggesting that IRE1α and IRE1β may differ at the level of their interactome. Thus, IRE1 BioID identified new IRE1 interactors and revealed that IRE1 interactions are responsive to stress.

### Protein-protein interactions involved in IRE1 signaling - The IRE1 signalosome

To parallel our approach with the IRE1 reference signalosome, we next analyzed the IRE1 BioID signalosome. We intersected the IRE1 BioID interactome with the results of the CRISPR and siRNA screens, as previously done with the reference interactome (see **Fig. 1D**). This identified 62 proteins constituting the IRE1 BioID signalosome (including IRE1 itself, **Fig. 4A, B**). Functional enrichment of this signalosome mainly revealed terms related to ER homeostasis (e.g., “protein targeting to ER”, “response to unfolded protein”, “protein folding in the ER”, “ER-Golgi transport”) (**Fig. 4B, C**). Comparison of the reference (**Fig. 1E**) and BioID (**Fig. 4A**) signalosomes revealed low overlap, with only 12 shared interactors (**Fig. S3A**). Moreover, enrichment in terms related to the IRE1 signalosome represented 30% of the BioID interactome compared to only 15% of the reference interactome, indicating an increased relevance (towards IRE1 signaling control) of the IRE1 interactome *in situ* (**Fig. S3B**). To evaluate the direct PPi between IRE1 and the components of the BioID signalosome, we used a deep learning-based algorithm (iPIN, see Materials and Methods section) and identified 50 predicted direct interactions in the whole BioID, of which 10 are part of the signalosome (**Fig. 4B & Table 1**).

**Figure 4:**
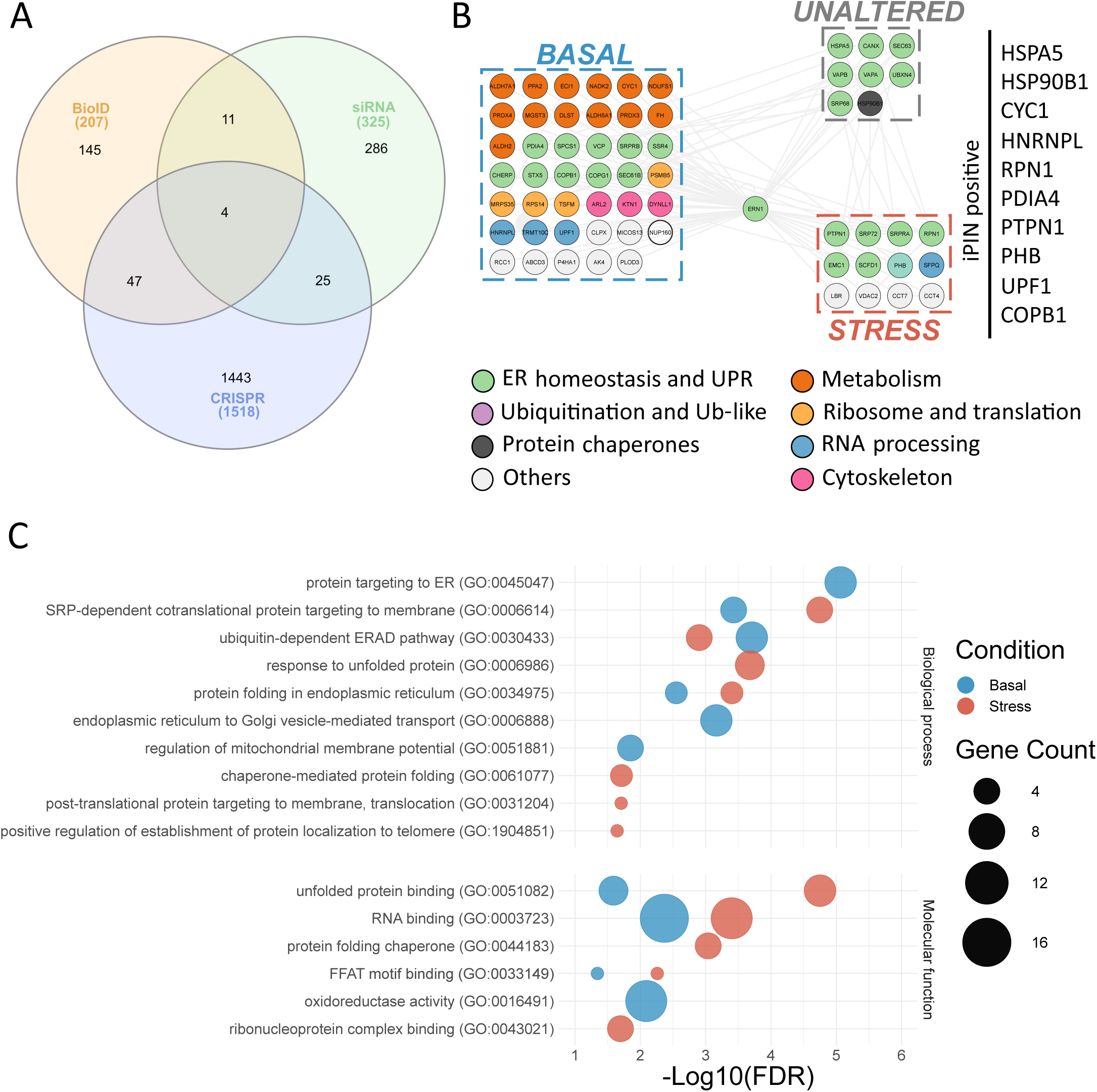
The BioID-based IRE1 signalosome. **A)** Venn diagram intersecting the lists of the IRE1 BioID interactome with siRNA and CRISPR screens and defining the IRE1 BioID signalosome. **B)** PPi network of the IRE1 BioID signalosome containing 62 proteins. Colors represent the functions associated after manual curation. In silico prediction of direct interactions using the iPIN algorithm retrieved 10 proteins predicted to directly interact with IRE1 and listed on the right. **C)** Comparison of Biological process (Top) and Molecular functions (bottom) Gene Ontologies of basal and stress IRE1 BioID signalosomes.

Our cytosolic BioID identified several ER lumen resident proteins (*HSP90B1, HSPA5,* and *PDIA4*) that should not be biotinylated, that are comprised in the signalosome (**Fig. 4B**). Surprisingly, these proteins were also predicted to interact with IRE1 cytosolic domain, which could be the case if subjected to reflux into the cytosol, as previously reported (Igbaria et al., 2019; Lajoie and Snapp, 2020; Sicari et al., 2021). Direct interactions with IRE1 were also predicted for the coat complex subunit *COPB1*, involved in retrograde transport, the phosphatase *PTPN1* (also known as *PTP1B)*, already known to regulate IRE1 signaling through RtcB (Gu et al., 2004; Papaioannou et al., 2022), *RPN1*, an Oligosaccharyl Transferase (OST) subunit, *HNRNPL* and *UPF1*, involved in mRNA maturation and RNA degradation, respectively, along with *PHB* and *CYC1*. All these IRE1 partners were also predicted to interact with each other, but not with RtcB (**Fig. S3C**). Among these interactions, only the interaction between IRE1 and COPB1 was confirmed in a yeast two-hybrid (Y2H) assay (**Fig. S3D**). Due to the limitations of Y2H, we pursued the characterization of specific signalosome components based on the positive iPIN hits COPB1, HRNRPL, PDIA4, PHB, PTPN1, RPN1 and UPF1.

### Regulation of canonical IRE1 signaling by components of the signalosome

To evaluate the impact of the aforementioned signalosome components on IRE1 signaling, we measured the effect of their silencing by siRNA knock down (KD) (**Fig. S4A**) on the different steps of the IRE1 signaling cascade, from oligomerization to subsequent RNase activities. IRE1 oligomerization was assessed by resolving samples on NativePAGE gels, which allows to distinguish the IRE1 monomer from a smear of higher order oligomers. Strikingly, the comparison between the monomer and oligomer bands shows increased oligomerization upon siCOPB1 and siPDIA4 (**Fig. 5A**). We then tested IRE1 RNase signaling. First, we measured *XBP1s* mRNA expression by RT-qPCR in unstressed cells. KD of *RtcB* was used as a positive control. This demonstrated that *HNRNPL*, *RPN1* and *PDIA4* KD resulted in decreased *XBP1s* expression (**Fig. 5B**). In contrast, siCOPB1 increased *XBP1* mRNA splicing (**Fig. 5B**). When treated with Tg, *PDIA4* and *PTPN1* KD led to a lower *XBP1* mRNA splicing induction (i.e. evaluating the specific effect of the KD on IRE1 splicing activity) compared to control (**Fig. 5C**). We next assessed the impact of the siRNAs on IRE1 RIDD induction by measuring *CD59* mRNA expression, a known substrate of this activity upon ER stress (Hollien et al., 2009). Unexpectedly, while CD59 mRNA is degraded by IRE1 in control condition, we observed that KD of *RtcB* and all of our candidates resulted in steady state *CD59* mRNA expression (**Fig. 5D**). This indicates that their expression is required for RIDD, however, this is hardly consistent with the results obtained with the oligomerization, suggesting different regulatory mechanism. Taken together, these RT-qPCR results suggests that our interactors of interest selectively regulate IRE1 RNase activity. In the native-PAGE experiments, we noted that overall IRE1 signal was not stable following siRNA KD, and observed a decrease in IRE1 expression in siHNRNPL transfected cells, while there is an increase of IRE1 expression in siUPF1 and siCOPB1 treated cells (**Fig. 5A)**.

**Figure 5:**
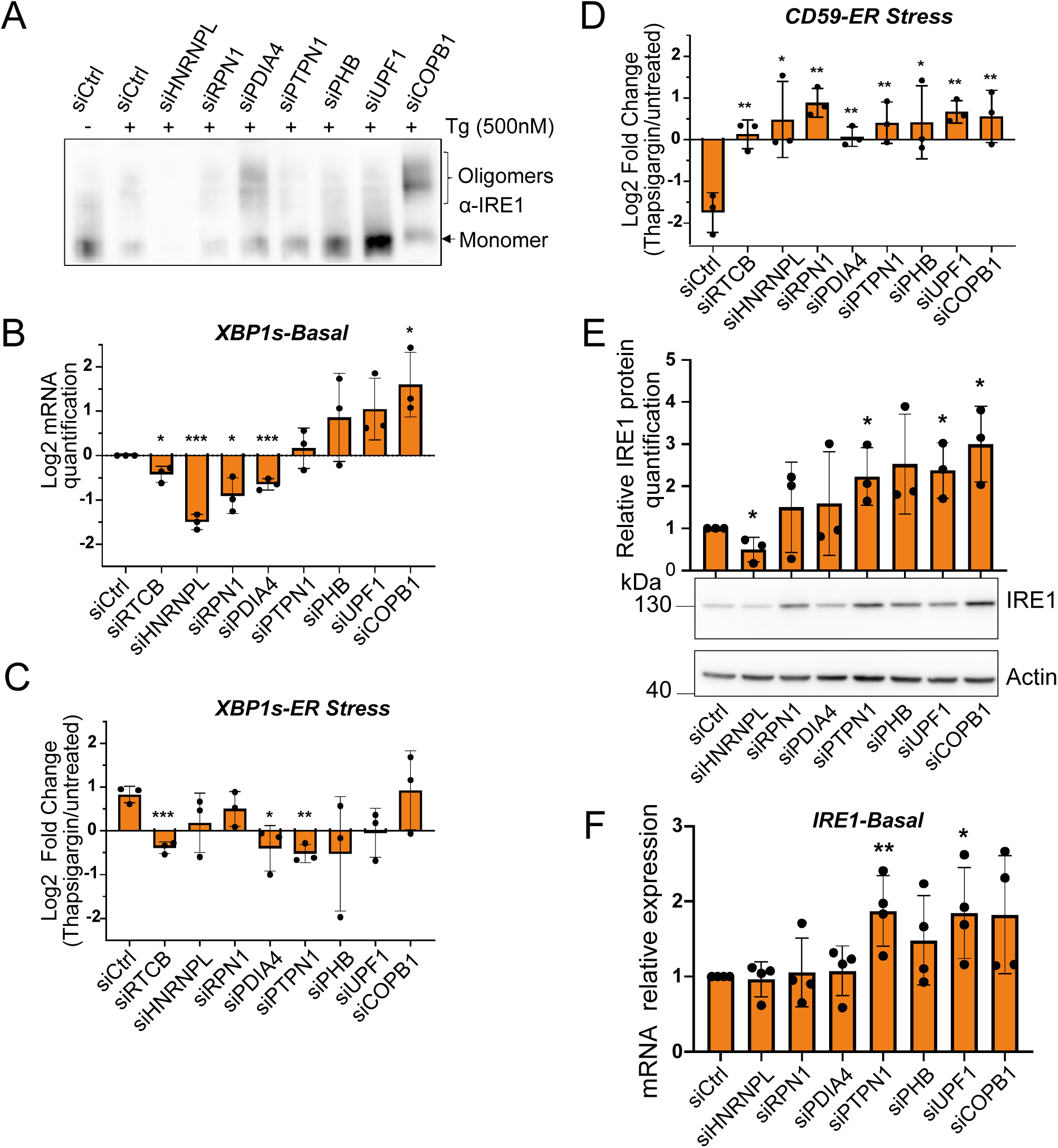
Components of the signalosome regulate IRE1 signaling. **A)** Western-Blot following separation by native gel electrophoresis of IRE1 oligomerization. HEK293T cells were transfected with 50 nM siRNAs targeting each of the genes of interest for 48 hours and further treated with 500 nM Thapsigargin for 6 hours. One representative experiment out of three is shown. **B, C, D)** RT-qPCR analysis of XBP1s (**B**, **C**) and CD59 (**D**) gene expression. HEK293T cells were transfected with 50 nM siRNAs targeting each of the genes of interest for 48 hours. Cells were pretreated with Actinomycin D for 1 hour and left untreated (**B**) or treated with 500 nM Thapsigargin for 6 hours (**C**, **D**). Data represent the gene expression fold changes between untreated and Thapsigargin treated cells, depicting the effect of siRNA treatment on IRE1 activation upon ER stress. Statistical significance represents comparison with control condition siCtrl. (n = 3 - *P < 0.05, **P < 0.01, ***P < 0.001). **E)** Western-Blot (bottom) and relative quantification (top) of IRE1 protein expression according to siRNA-mediated silencing of genes of interest. One representative experiment out of three is shown. **F)** RT-qPCR analysis of IRE1 gene expression in HEK293T cells transfected with 50 nM siRNAs targeting each of the genes of interest for 48 hours. (n = 4 - *P < 0.05, **P < 0.01, ***P < 0.001).

This led us to investigate whether IRE1 expression was modulated by our proteins of interest. Evaluation of the protein expression levels by standard SDS-PAGE confirmed decreased IRE1 expression under siHNRNPL condition (**Fig. 5E**). In addition, we found that IRE1 protein expression was increased following *PTPN1*, *UPF1* and *COPB1* downregulation (**Fig. 5E**). For UPF1 and PTPN1 KD, increased IRE1 protein level could result from altered IRE1 RNA levels (**Fig. 5G**), whilst KD of HNRNPL and COPB1 did not alter IRE1 mRNA levels (**Fig. 5F**). This suggests that the downregulation of these proteins affects IRE1 at the protein level, and therefore may involve PPi. Of note, we analyzed the effect of these siRNA on the activation of the three UPR branches. We found that downregulation of *HNRNPL*, *PTPN1*, *PHB*, *COPB1* promotes the expression of *ERDJ4* mRNA, a target gene of XBP1s (**Fig. S4B**) (Lee et al., 2003). In addition, *HNRNPL*, *UPF1* and *COPB1* KD caused an increase in *CHOP* expression resulting from PERK activation (**Fig. S4C**) (Marciniak et al., 2004). Finally, the ATF6 branch was the least affected, as only silencing of RPN1 and PTPN1 resulted in increased expression of *SEL1L*, an ATF6 target gene (**Fig. S4D**) (Kroeger et al., 2018). However, *HERPUD1*, another ATF6 target gene, was not altered by siRNAs treatment (**Fig. S4E**). This indicates that the identified candidates could differentially fine-tune the intensity of each UPR branch (**Fig. S4F**).

### PTPN1 directly interacts with IRE1 to regulate the splicing of *XBP1*

Based on our previous studies describing IRE1 signaling regulation by PTPN1 (Gu et al., 2004) through the dephosphorylation of phosphotyrosine residues on RtcB (Papaioannou et al., 2022) and our current observations from the iPIN prediction that PTPN1 could directly interact with IRE1 and that its targeting impairs IRE1 signaling, we hypothesized that PTPN1 could regulate IRE1 through direct PPi. To test this, the complex was modeled by additional protein docking to the structure defined by Papaioannou *et al*. Here, PTPN1 was added to the complex formed by IRE1 tetramer and *XBP1* mRNA, and we were able to build a model where some IRE1 residues, namely N683, K748 and N750, bind to PTPN1 (**Fig. 6A**). We then tested whether mutations of these binding residues could alter the IRE1-PTPN1 complex. To do this, we calculated the free energy of binding of each contact site and observed that mutation of IRE1 683^th^ residue into an asparagine was the most altering modification (**Fig. 6B**). Hence, we designed and synthesized plasmids of a Flagged WT and IRE1 N683D mutant (**Fig. 6C**), whose protein expression were confirmed by Western blot after transient transfection in MA2 *IRE1* KO cells (**Fig. 6D**). Notably, IRE1 N683D appeared to be expressed at lower levels than WT. Next, we monitored the interaction of WT and N683D IRE1 with PTPN1 using co-immunoprecipitation upon overexpression of both IRE1 constructs and PTPN1 upon stress since the IRE1/PTPN1 interaction was observed under ER stress in the BioID experiment (**Fig. 3**) and both proteins were found to coimmunoprecipitate upon ER stress induction (**Fig. 6E**). Moreover, quantification of PTPN1 mCherry signal in the immunoprecipitates increased in cells overexpressing IRE1 N683D compared to those overexpressing IRE1 WT (**Fig. 6F**). This suggests an enhanced complex formation with PTPN1 upon modification of the IRE1-PTPN1 interface (**Fig. 6A, E, F**), thereby suggesting a stabilized interaction. To evaluate how this mutant affects IRE1 signaling, HEK293T cells were transfected with WT and N683D IRE1 constructs, treated with Tg for up to 12 hours, and *XBP1s* mRNA expression was monitored by qPCR. Despite its lower expression, IRE1 N683D exhibited higher *XBP1* mRNA splicing activity over time compared to WT (**Fig. 6G**), which may be due to more effective formation of the splicing complex.

**Figure 6:**
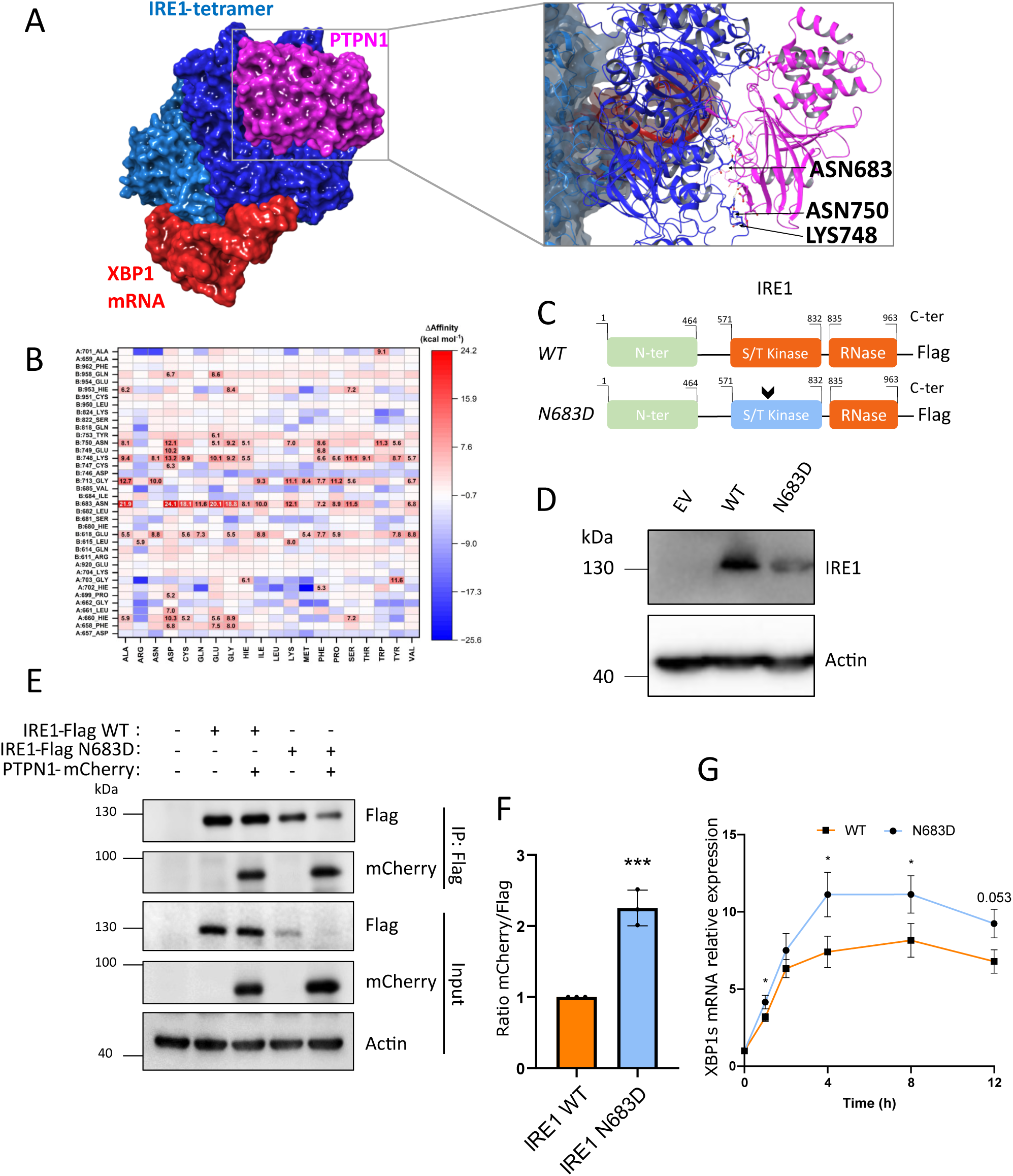
PTPN1 directly interacts with IRE1 to regulate the splicing of XBP1. **A)** Docked complex of IRE1 tetramer/XBP1 mRNA/PTPN1. **B)** Heat-map representing the relative free energy of binding of the interaction interfaces between IRE1 and PTPN1 according to IRE1 mutations by other amino acids. **C)** Schematic representation of IRE1 WT and IRE1 N683 mutant disrupting IRE1-PTPN1 interaction. The arrow represents the approximative location of the mutation. **D)** Western-Blot of IRE1 wild-type and mutant transiently expressed in MA2 IRE1 KO cells. One representative experiment out of three is shown. **E)** HEK293T cells were transfected with empty vector (EV), IRE1 WT and N638D (each containing a Flag tag) with or without PTPN1-mCherry and treated with 500 nM Thapsigargin for 4h. IP was performed in the cell lysates using Flag Ab and the immunoprecipitates were immunoblotted for PTPN1 and Flag. Input samples probed for PTPN1 and Flag. One representative experiment out of three is shown. **F)** Quantification of the association between IRE1 and PTPN1 assessed by calculating the ratio between PTPN1 mCherry and IRE1 Flag signals in the immunoprecipitates (n = 3 - *P < 0.05, **P < 0.01, ***P < 0.001). **G)** RT-qPCR analysis of XBP1s gene expression in HEK293T cells transfected with IRE1 wild-type or IRE1 N683D plasmids. After 48 hours transfection, cells were treated with 500 nM Tg for 0, 1, 2, 4, 8, 12 hours. (n = 3 - *P < 0.05, **P < 0.01, ***P < 0.001)

### HNRNPL protects IRE1 from ER associated degradation

In our previous experiments, we observed that IRE1 protein expression was reduced upon HNRNPL siRNA-mediated silencing (**Fig. 5E**) and that this was independent of *IRE1* mRNA expression (**Fig. 5F**). This observation was confirmed in various cell lines (**Fig. S5**). IRE1 protein expression analysis using Western blot revealed that HNRNPL silencing resulted in decreased IRE1 expression and that this was rescued upon proteasome inhibition with MG132 under non-stressed conditions (**Fig. 7A, B**). In line with our results, a study previously showed that IRE1 is a client for ERAD (Sun et al., 2015). We therefore tested whether HNRNPL could contribute to ERAD-mediated IRE1 degradation under basal conditions. To this end, we co-transfected cells with siRNAs targeting *HNRNPL* and *SYVN1*, a major ubiquitin ligase involved in ERAD (Bays et al., 2001). Remarkably, downregulation of *SYVN1* rescued IRE1 expression attenuation caused by HNRNPL deficiency (**Fig 7C, D**). At last, we evaluated the expression levels of IRE1 over time under cycloheximide treatments, in cells silenced or not for HNRNPL, and under basal or Tg-induced ER stress. These experiments revealed that IRE1 protein half-life was dramatically shortened upon HNRNPL silencing under basal conditions (**Fig. 7E, G**) but not under stress (**Fig. 7F, H**). These observations are the first to report the regulation of IRE1 protein expression by HNRNPL and the links to ERAD.

**Figure 7:**
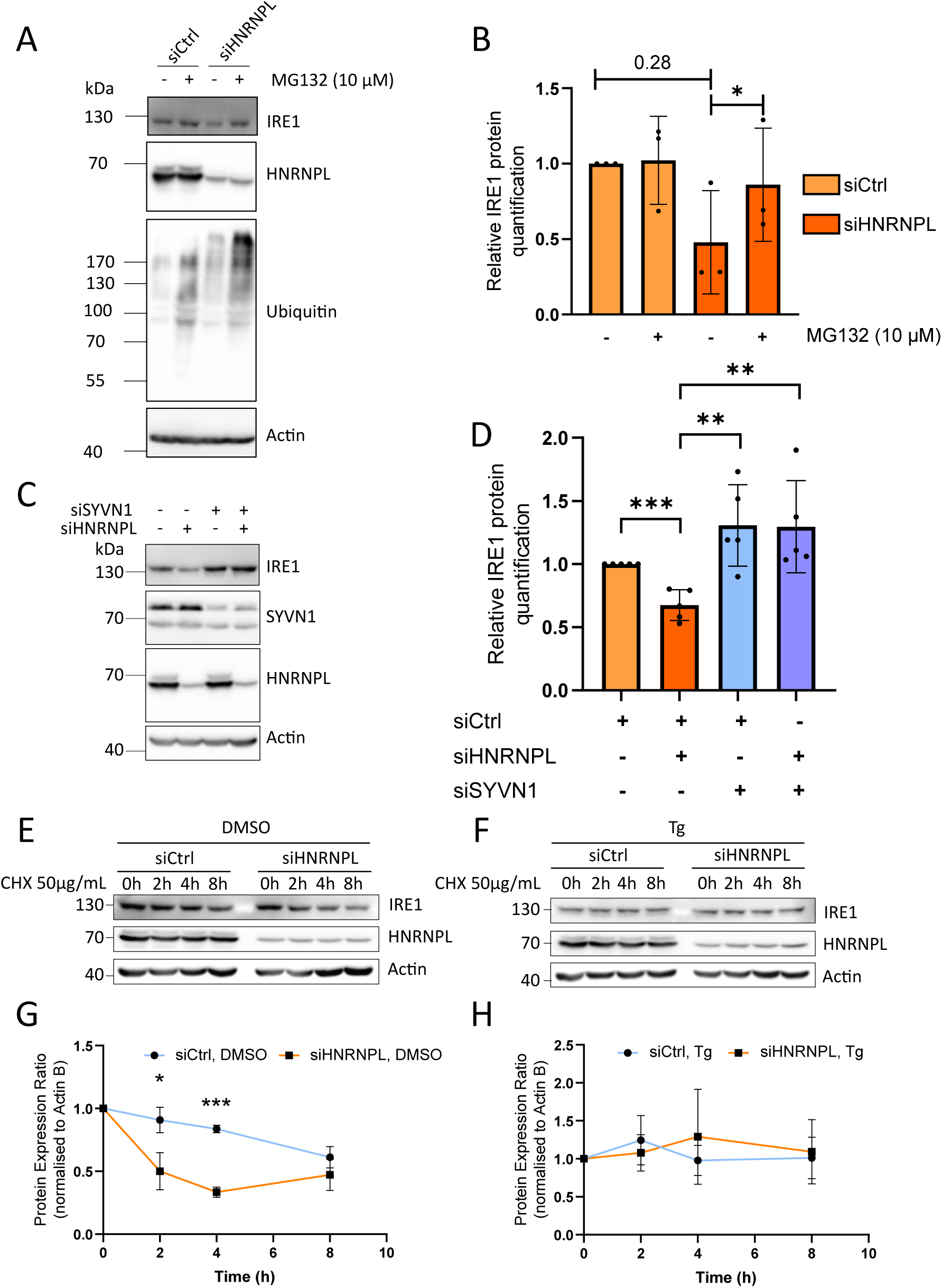
HNRNPL protects IRE1 from ER associated degradation. **A)** IRE1 protein expression assessed by Western blot according to HNRNPL downregulation by 50 nM siRNA for 48 hours and rescue of the siHNRNPL phenotype with the proteasome inhibitor treatment MG132 for 8 hours. One representative experiment out of three is shown. **B)** Quantification of A. (n = 3 - *P < 0.05, **P < 0.01, ***P < 0.001)**. C)** IRE1 protein expression assessed by Western blot according to HNRNPL and SYVN1 downregulation by 50 nM siRNA for 48h. One representative experiment out of four is shown. **D)** Quantification of C (n = 4 - *P < 0.05, **P < 0.01, ***P < 0.001). **E, F)** Western blot analysis of IRE1 expression upon cycloheximide (CHX) exposure, and siCtrl or siHNRNPL treatment, and under basal (**E**) or 500 nM thapsigargin-induced ER stress conditions (**F**). One representative experiment out of three is shown. **G, H)** Quantification of E and F, respectively (n = 3 - *P < 0.05, **P < 0.01, ***P < 0.001).

## Discussion

In this study, we further document the IRE1 signalosome as previously defined (Le Goupil et al., 2024) by combining *in vitro* and *in silico* analyses, and proposing a new method to assess whether a PPi impair IRE1 signaling (**Fig. 1 & 4**). We found that a subset of proteins from diverse cellular pathways are able to fine-tune IRE1 signaling (comprising *XBP1* mRNA splicing and RIDD) and specifically, that most of them are required for *CD59* mRNA cleavage through RIDD activity. This highlights the interplay between the UPR and the other proteostasis-related processes. In particular, our results are consistent with the close association between IRE1 and the co-translocation translation machinery (Acosta-Alvear et al., 2018), while uncovering close relationships with COP transport (COPB1) and other RNA metabolic pathways such as their maturation or degradation by Nonsense mediated Decay (NMD). NMD has been shown to occur near the ER (Longman et al., 2020), and we observed that UPF1 is important for ensuring IRE1 RIDD activity (**Fig. 5**). UPF1 hyperphosphorylation (on serine and threonine) promotes NMD (Durand et al., 2016), and *in silico* prediction of UPF1 phosphorylation (phosphosite plus) shows a high prediction score for an IRE1-dependent phosphorylation at S10, which may be in line with the increasing number of proteins found to be phosphorylated by IRE1 (Cairrão et al., 2022; A. D. Yildirim et al., 2022; Z. Yildirim et al., 2022). Taken together, we can hypothesize that RIDD and NMD are involved in a crosstalk to ensure proper degradation of mRNA aiming to decrease the translational load. We further propose that this interaction could occur at ER contact sites with membrane-less organelles, structures highly enriched in RNA and RNA binding proteins (RNP granules), such as stress granules. This is supported by the identification of G3BP1 and UPF1 in our BioID, proteins involved in RNP granules biogenesis and NMD, respectively (Fischer et al., 2020; Kedersha et al., 2016). In addition, RNP granules localize near the ER (Child et al., 2021). Hence, whether IRE1 localize at ER-targeted RNP granules should be taken into consideration for further study.

In parallel, we revealed that another RNA binding protein whose canonical roles are related to RNA transport and maturation, the ribonucleoprotein HNRNPL is important for IRE1 stability (**Fig. 5 & 7**). Futur studies will focus on elucidating the mechanisms involved in such stabilization. Here we propose HNRNPL as a novel regulator of the UPR, serving to transport RNA substrates to IRE1 for degradation while also stabilizing IRE1 under basal conditions, thereby protecting it from proteasomal degradation. While IRE1 is known to be an ERAD substrate (Sun et al., 2015), preliminary results from our lab suggest that IRE1 may be degraded by an alternative proteasomal-dependent mechanism (**Fig. 7**). Since we and others have identified several proteins involved in ERAD in our BioID, including VCP and UBXN4 (Ahmed et al., 2024; Liang et al., 2006), we hypothesize that there is a competing interaction between HNRNPL and the ERAD machinery (i.e. VCP, UBXN4, SYVN1, SEL1L) in regulating IRE1 protein expression. Given that several HNRNPs including HNRNPL have been observed in RNP granules in the cytoplasm of HEK293T or HeLa cells under basal conditions (Guil et al., 2006; J⊘nson et al., 2007; Wall et al., 2020) and that IRE1 coalesces with stress granules upon ER stress (Liu et al., 2024), one could hypothesize that HNRNPL-granules associate with IRE1 at basal levels, physically protecting IRE1 from ubiquitination by SYNV1 and subsequent degradation. Upon stress, HNRNPL granules are switched for stress granules. Therefore, IRE1 stability becomes independent of HNRNPL granules. At last, the phosphotyrosine phosphatase PTPN1 has been identified to be involved in IRE1 signaling and was later described to dephosphorylate RtcB, leading to increased *XBP1* mRNA splicing (Gu et al., 2004; Papaioannou et al., 2022). Here, we provide the first evidence that PTPN1 is in a complex with IRE1. Functional assays with IRE1 mutant showed that stabilization of IRE1-PTPN1 interaction leads to increased *XBP1* mRNA splicing activity. Therefore, there may be a synergistic effect where PTPN1 interacts with IRE1 prior to RtcB to act as a recruiting (and stabilizing) platform for the IRE1-RtcB-PTPN1 complex thereby favoring *XBP1* mRNA splicing.

### Approach limitations

Aside from the canonical roles associated with the management of ER stress, IRE1 has been characterized to participate in other cellular processes. These non-canonical roles are highly dependent on PPi (Hetz et al., 2020). Here, we identified new IRE1 interactors using a BioID-MS approach. It should be noted that the 18-24h treatment of the BioID labeling does not provide a snapshot of the PPis at a specific time, but the whole of the interactions that happened during the labeling. Concomitantly with a recent study combining TurboID with IRE1, another BioID-based approach with faster enzymatic activity (Ahmed et al., 2024), we provide an *in situ* IRE1 interactome, with physiologically relevant interactions (**Fig. 3**). In line with the respective techniques, our BioID study shared few interactors with the reference interactome. However, our interactome identified previously described IRE1 interactors such as the actin-binding protein FLNA (Urra et al., 2018), the Ca^2+^ influx regulator STIM1 (Carreras-Sureda et al., 2023), Sec61 translocon subunits (Sundaram et al., 2017), as well as VCP and MTDH, which were also covered by the TurboID study (Ahmed et al., 2024). This therefore confirms the relevance of such an approach. While we share common observations with Ahmed and colleagues, the different kinetics of the two BioIDs might challenge the temporality of IRE1 interactions, as TurboID exerts faster biotinylation compared to BioID (Guo et al., 2023). Since we used transient overexpression of IRE1 for our BioID study, there might be an increased basal level of ER stress compared to stable transfection, modifying the basal UPR signaling properties. In addition, it might be possible that minor amounts of IRE1 may result in interaction with the ERAD machinery or chaperones, a possibility that requires further investigation.

Overall, our study reveals an integrative perspective in the management of ER homeostasis by linking its various aspects (e.g. trafficking, translation, protein expression) to IRE1 signaling. In addition, we highlight new mechanisms regulating IRE1 signaling and their ability to fine-tune ER homeostasis.

## Materials and Methods

### Cell culture and transfection

Human HEK293T, A375-MA2, HeLa, MDA-MB-231, SUM 159, U87 and U251 cells were cultured in Dulbecco Modified Eagle Medium (DMEM) (Gibco, 41966-029) supplemented with 10% fetal bovine serum (FBS) (Gibco, 10437-028). Cells were maintained in a humidified incubator at 37°C and 5% CO_2_. Cells were transiently transfected with pcDNA3.1-IRE1-BirA*, pcDNA3.1-BirA*, pTwist-IRE1 Wild-type (Twist Bioscience), pTwist-IRE1 N683D (Twist Bioscience) and pPTP1BD181A-mCherry (gift from Martin Offterdinger, Addgene plasmid #40270 (Offterdinger and Bastiaens, 2008)) using either Polyethylenimine (PEI; Sigma Aldrich, 408727) for HEK293T and A375-MA2 cells or Lipofectamine 2000 (ThermoFisher Scientific, 11668019) for HeLa cells. The siRNAs were obtained from Dharmacon, as siGENOME SmartPool for targeted mRNA and non-targeting mRNA (**Table S1**). Each siRNA (50 nM) was transfected by reverse transfection using Lipofectamine RNAiMAX (ThermoFisher Scientific, 13778075) according to manufacturer’s instruction. ER stress was induced using thapsigargin (50 or 500 nM according to specifications) or tunicamycin (5 µg/ml). When measuring XBP1 splicing and RIDD activities by RT-qPCR, cells were preincubated with Actinomycin D (ThermoFischer Scientific, A7592) for 1 hour before further treatment.

### Generation of IRE1 knock out MA2 cell line

A375-MA2 IRE1 knock out cells were generated using the double nickase method of CRISPR/CAS9 technology. To this end, a double nickase targeting either IRE1 or a scrambled control (sc-400576-NIC and sc-437281; Santa Cruz) was used. A375-MA2 cells were transfected using an Effectene protocol (301425, Qiagen). Transfected cells were subsequently selected with 2 µg/ml of puromycin. Individual clones derived from limiting dilutions were isolated and subsequently expanded. Clones were expanded and selected based on their IRE1 expression and activity.

### Western-Blot and immunoprecipitation

Cells were harvested and lysed using RIPA buffer (50 mM Tris-HCl pH 7.5, 150 mM NaCl, 1 mM EDTA, 0.2% SDS, 1% Triton X-100, 1% sodium deoxycholate) containing proteases (Merck, 4693159001) and phosphatases inhibitors (Merck, 4906845001). Samples are sonicated and centrifugated (15,000g, 15 minutes) after which protein-containing supernatant were collected. Protein concentration was determined using BCA protein assay kit (Pierce, 23227). 20-40 µg of protein were separated on SDS-PAGE, transferred onto nitrocellulose membrane, and blocked with 5% bovine serum albumin in Tris-buffered saline/Tween 20 (TBST) at room temperature. Membranes were incubated at 4°C overnight with the corresponding primary antibody, which had been diluted in TBST with 5% BSA to the appropriate concentration. Membranes were then washed three times in TBST. Secondary HRP-coupled antibodies diluted in TBST with 5% BSA were applied to the membranes for 1 hour at room temperature, followed by three washes with TBST. Detection of the signal was conducted using ECL RevealBlot Intense (Ozyme, OZYB002-1000) in a Gbox Chemi XX6 imager (Syngene). Antibodies used in this work are listed in **Table S2**. To evaluate IRE1 oligomerization, NativePAGE (Thermo Fischer Scientific, BN1001 BOX) was conducted following non-denaturing protein extraction with digitonin. Extracted proteins were then transferred onto a PVDF membrane. For the detection of IRE1 phosphorylation, PhosTag^TM^ (Sobioda, W1W304-93526) was used following manufacturer’s instruction, with the aforementioned protein extraction method (SDS-PAGE). For the IP, the cells were lysed in lysis buffer (50 mM Tris–HCl pH 7.4, 150 mM NaCl, and 1% CHAPS) supplemented with a cocktail of protease inhibitors. The lysates were incubated for 16 hours at 4°C with the indicated IP antibody (1 μg Ab per 1,000 μg protein). Subsequently, dynabeads protein G (Thermo Fisher Scientific, 88803) were washed with lysis buffer, then mixed with the protein/Ab mixture. This was incubated at 4°C for 1 hour with gentle rotation. The protein/Ab/beads mixture was washed with lysis buffer, and beads were eluted with 5X Laemmli sample buffer, heated at 95°C for 5 minutes. The mixture was then loaded onto an SDS–PAGE. For immunoblotting, suitable primary and secondary antibodies were used.

### Immunofluorescence

Cells were plated on cover slips within 12 well plates, transfected and treated with biotin as described above. Cells were washed four times with PBS and fixed with 4% formaldehyde in PBS for 15 minutes at room temperature. Subsequently, cells were washed four times with PBS and treated with permeabilization buffer (PBS, 3% BSA, 0.2% Triton X-100) for 20 minutes at room temperature, then washed again four times with PBS. Cells were incubated at 4°C overnight with the corresponding antibody, which had been diluted in PBS with 3% BSA. They were then washed four times with PBS. Secondary antibody was applied for 3 hours at room temperature, after which cells were washed four times with PBS. Cover slips were positioned on a microscope slide and sealed with DAPI mounting medium (Sigma-Aldrich; DUO82040). Slides were observed with a Leica SP8 confocal microscope using a PLAN APO x63/1.40 oil objective.

### RNA extraction and RT-qPCR

Total RNA was extracted using TRIzol (Thermo Fischer Scientific, 15596018) according to manufacturer’s instructions. A total of 2 µg of RNA was reverse transcribed using Maxima Reverse Transcriptase (Thermo Fischer Scientific, EP0743) and random hexamer primers (Thermo Fischer Scientific, SO412) according to manufacturer’s instructions. Quantitative PCR was conducted using SYBR Premix Ex Taq (TAKARA Bio, RR420L) on a QuantStudio5 system (Applied Biosystems) with the appropriate primers (**Table S3**). The levels of target transcripts were normalized with GAPDH.

### BioID, biotinylation and mass spectrometry analysis

This BioID method was adapted from the Soderling lab (Uezu et al., 2016). In brief, cells transfected with IRE1-BirA* and BirA* were treated with 50 µM biotin for 18 to 24 hours. Cells were lysed with RIPA buffer (50 mM Tris-HCl pH 7.5, 150 mM NaCl, 1 mM EDTA, 0.2% SDS, 1% Triton X-100, 1% sodium deoxycholate) supplemented with protease inhibitors, and lysates were sonicated. For the streptavidin pulldown, 20 µL of slurry neutravidin-coupled beads was added to the lysate, which was then incubated at 4°C overnight. On the subsequent day, beads were washed consecutively as follows: two washes with 2% SDS, two washes with the following buffer (1% Triton X-100, 1% sodium deoxycholate, 25 mM LiCl), two washes with 1 M NaCl, and two washes with 50 mM Ambic. Elution was conducted by heating samples at 95°C with 5X sample buffer complemented with 1 mM biotin. Mass spectrometry analysis was conducted using the same protocol, with the exception of the elution step. Here trypsin digestion on bead (3 hours) was conducted in 96 microplates following reduction with 10 mM DTT and alkylation with 20 mM Iodoacetamide, both for 30 min. Samples were acidified with 0.5% formic acid and supernatants were transferred to adjacent wells for LC-MS. Mass spectrometry was conducted using an Ultimate 3000 nano-RSLC coupled in-line with an Exploris 480 orbitrap via a nano-electrospray ionization source (Thermo Scientific, San Jose California) and the FAIMS pro interface. Samples (1µl) were loaded onto a C18 Acclaim PepMap100 trap-column (75 µm ID x 2 cm, 3 µm, 100Å, Thermo Fisher Scientific) for 3.5 minutes at 6 µL/min with 2% ACN, 0.1% FA in H_2_O. Samples were then separated on a C18 PepMap nano-column (75 µm ID x 15 cm, 2.6 µm, 150 Å, Thermo Fisher Scientific) with a 45-minute linear gradient from 7% to 35% buffer B (A: 0.1% FA in H_2_O; B: 0.1% FA in 80% ACN, 350 nl/min, 45°C). This was followed by a regeneration step at 90% B and an equilibration step at 7% B. The total chromatography time was 60 minutes. The mass spectrometer was operated in positive ionization mode in Data-Dependent Acquisition (DDA) with FAIMS compensation voltages set up −45V. The DDA cycle consisted of one survey scan (350-1400 m/z, 120,000 FWHM) followed by MS² spectra (HCD; 30% normalized energy; 2 m/z window; 30,000 FWMH) within the limit of 1 second. The Ion Target Value for the MS1 scan and the MS2 scan were set to 3E6 and 1E5, respectively, and the maximum injection time was set to 50 ms for both scan modes. Unassigned and single charged states were excluded. Exclusion duration was set for 40 seconds with mass width was ± 10 ppm. Proteins were identified with Proteome Discoverer 2.5 software (Thermo Fisher Scientific) and the Homo sapiens proteome database (https://www.uniprot.org/proteomes/UP000005640). Precursor and fragment mass tolerances were set at 7 ppm and 0.05 Da, respectively, and up to 2 missed cleavages were allowed. Oxidation (M, +15.994) was set as variable modification, and Carbamidomethylation (C) as fixed modification. Peptides were filtered with a false discovery rate (FDR) of 1%, rank 1. Proteins were quantified with a minimum of 1 unique peptide based on the XIC (sum of the Extracted Ion Chromatogram). The resulting quantification values were exported in Perseus 1.6.15.0 for log[2] transformation, imputation, normalization to the median and statistical analysis (Tyanova et al., 2016).

### Modelling of protein complexes

Interactions between IRE1 and PTPN1 phosphatase were studied using a recently developed model of the human IRE1-tetramer complexed with XBP1-mRNA, both with and without RtcB ligase (Papaioannou et al., 2022), and the crystal structure of the human PTPN1 phosphatase (PDB-id: 8SKL, 1.5 Å) (Liang et al., 2023). Protein-protein docking was conducted using eight different protein docking engines: Cluspro (Kozakov et al., 2017), GalaxyWEB (Ko et al., 2012), GRAMM-X (Tovchigrechko, Andrey and Vasker, Ilya, 2006), HDOCK (Yan et al., 2020), LZerD (Christoffer et al., 2021), PyDockWEB (Jiménez-García et al., 2013), ZDOCK (Chen et al., 2003), and MOE (Chemical Computing Group. 2023. www.chemcomp.com). This approach was based on the meta-approach outlined by Mahdizadeh et al. (Mahdizadeh et al., 2021). For each engine, the top 10 predicted complexes were selected using the default settings and parameters. Subsequently, each of the 80 initial docking results was refined using the GalaxyRefineComplex tool under two distinct relaxation protocols (Heo et al., 2016). The first protocol applied only distance restraints, while the second applied both distance and positional restraints. The five lowest energy complexes from each protocol were selected, resulting in 10 refined models for each initial complex. These 800 refined models were then clustered based on RMSD values of all heavy atoms using the “clustering of conformer” module in Maestro Schrödinger (Schrödinger Release 2023-4: Maestro, Schrödinger, LLC, New York, NY, 2023). The optimal number of clusters was determined using Kelley penalty plots (Kelley et al., 1996). The model closest to the centroid of the most populated cluster was chosen as the final model. To identify the key residue involved in the interaction between IRE1 and PTPN1, all interfacial amino acids of IRE1-tetramer within a cut-off distance of 5 Å from PTPN1 were mutated to other 19 standard amino acids. The impact of each mutation on the variation of free energy of binding was then calculated using the thermodynamic cycle through the molecular mechanics generalized Born surface area (MM-GBSA) technique (Rastelli et al., 2010) with OPLS4 force field (Lu et al., 2021).

### Y2H interaction assays and associated cloning

The coding sequence of IRE1 K599A was PCR-amplified and cloned in frame with the Gal4 DNA binding domain (DBD) as a C-terminal fusion to Gal4 (Gal4-bait fusion) into the pAS2ΔΔ-derived vector pB66 plasmid (Fromont-Racine et al., 1997). The coding sequences of the HNRNPL, RPN1, PDIA4, PTPN1, UPF1, and COPB1 proteins were cloned in frame with the Gal4 Activation Domain (AD) into plasmid pP7 (AD-prey). Additionally, PTPN1 was cloned as an N-terminal fusion to the Gal4 Activation Domain (AD) into plasmid pP13 (prey-AD). The two prey plasmids were derived from the original pGADGH (Bartel et al., 1993). The bait and prey constructs were transformed into the yeast haploid cells CG1945 (matα) and YHGX13 (Y187 ade2-101::loxP-kanMX-loxP, matα), respectively. The diploid yeast cells were obtained through a mating protocol with both yeast strains. The assays are based on the HIS3 reporter gene, which is used to assess growth in the absence of histidine. As negative controls, the bait plasmid was tested in the presence of empty prey vector (pP7/pP13) and all prey plasmids were tested with an empty bait vector (pB66). The interaction between SMAD and SMURF is used as positive control (Colland et al., 2004). Controls and interactions were tested in the form of streaks of three independent yeast clones for each control and interaction on DO-2 and DO-3 selective media. The DO-2 selective medium lacking tryptophan and leucine was used as a growth control and to confirm the presence of the bait and prey plasmids. The DO-3 selective medium, which lacks tryptophan, leucine, and histidine is used to select for the interaction between the bait and prey.

### Bioinformatics tools and analyses

To generate the reference IRE1 interactome, data from the following public repositories were retrieved: Biogrid, IntAct, String, and Agile Protein Interactomes DataServer (APID) (Alonso-López et al., 2019; Orchard et al., 2014; Oughtred et al., 2021; Szklarczyk et al., 2019). These data were pooled with data extracted from a study using native immunoprecipitation and mass spectrometry with IRE1 as bait in human cells (Acosta-Alvear et al., 2018). The Acosta-Alvear dataset was curated by selecting the 400 proteins with the highest interaction specificity score, defined as the normalized number of unique peptides described by the authors [number_of_unique_peptides/142) x 2] and weighted by the protein contribution to the first principal component of a PCA performed on LFQ intensities. Protein-protein interaction networks were built using the Cytoscape software (Shannon et al., 2003) (https://cytoscape.org, Version 3.10.0) with the standard confidence score threshold of > 0.4 and based on the STRING database (https://string-db.org, Version 11.0), which is designed to assess protein physical or functional associations (Szklarczyk et al., 2019). Gene Ontology enrichment analysis was conducted using the PANTHERdb to retrieve functional enrichment for GO Biological Process and GO Molecular Function (Thomas et al., 2022). Redundant terms were manually removed, and the top enriched terms were displayed. Subcellular compartmentalization of the proteins was assessed using the COMPARTMENTS plugin in Cytoscape, with a cut-off of 4.75 (95%) (Binder et al., 2014). iPIN (intelligence Protein Interaction Network) is an AI-based protein interaction prediction tool developed by S. J. Mahdizadeh at university of Gothenburg, Sweden, and ANYO Labs AB (www.anyolabs.com). iPIN is capable of predicting interacting proteins from their amino acid sequence information, quickly and accurately. iPIN was trained using ∼70,000 protein pairs from the human proteome (50/50 positive and negative pairs collected from the Human Protein Reference Dataset (HPRD) 2009 and Swiss-Prot 57.3, respectively). The performance of iPIN was extensively benchmarked against interactome data available for a diverse range of eukaryotic (*H. sapiens, M. musculus, C. elegans, D. melanogaster*, *S. cerevisiae*) and prokaryotic (*H. pylori* and *E. coli*) organisms. Venn diagrams were constructed using the InteractiVenn tool (http://www.interactivenn.net/index2.htm).

### Statistical analyses

Data are expressed as means ± SD of at least three independent experiments. P values were analyzed using Student’s *t*-test or ANOVA using Prism 8 (GraphPad). *χ²* test was performed with R studio (Build 386).

## Supporting information

LeGoupil_et_al_Suppl

## Data availability

The mass spectrometry proteomics data resulting from IRE1 BioID have been deposited to the Proteome Xchange via the PRIDE partner repository and MassIVE Consortiums with the dataset identifiers PXD051657 and MSV000094379 respectively. Data are available at: (https://proteomecentral.proteomexchange.org/ui) and (https://massive.ucsd.edu/ProteoSAFe/static/massive.jsp).

## Acknowledgements

We thank Dr E. Lafont for critical reading of the manuscript. This work was funded by grants from Institut National du Cancer (INCa, PLBio2019, 2020, 2022) and Fondation pour la Recherche Médicale (FRM EQU202403018041) to EC. SLG was funded by a PhD scholarship from the French Ministry of research, HL was funded by a post-doctoral fellowship from Région Bretagne, CP was funded by a Marie Sklodvska Curie Action Bienvenü program from the European Commission to the Brittany region.

## Authors contribution

SLG: conceptualization, formal analysis, investigation, visualization, methodology, and writing—original draft; HL: conceptualization, formal analysis, investigation, visualization, methodology; ML: visualization; RB: investigation; DPR: investigation; CP: investigation; KD: investigation and formal analysis; LN: mass spectrometry experiment and data analysis; KD: mass spectrometry experiment and data analysis; GJ: mass spectrometry experiment and data analysis; CL: Resources; CH: resources, and funding acquisition; JM: investigation, formal analysis and methodology; JER: conceptualization and funding acquisition; LE: investigation, formal analysis, funding acquisition, visualization, methodology; E Chevet: conceptualization, formal analysis, supervision, funding acquisition, validation, investigation, project administration, and writing—original draft, review, and editing.

## References

Acosta-Alvear, D., Karagöz, G.E., Fröhlich, F., Li, H., Walther, T.C., Walter, P., 2018. The unfolded protein response and endoplasmic reticulum protein targeting machineries converge on the stress sensor IRE1. eLife 7, e43036. 10.7554/eLife.43036

Adamson, B., Norman, T.M., Jost, M., Cho, M.Y., Nuñez, J.K., Chen, Y., Villalta, J.E., Gilbert, L.A., Horlbeck, M.A., Hein, M.Y., Pak, R.A., Gray, A.N., Gross, C.A., Dixit, A., Parnas, O., Regev, A., Weissman, J.S., 2016. A multiplexed single-cell CRISPR screening platform enables systematic dissection of the unfolded protein response. Cell 167, 1867–1882.e21. 10.1016/j.cell.2016.11.048

Ahmed, N., Preisinger, C., Wilhelm, T., Huber, M., 2024. TurboID-Based IRE1 Interactome Reveals Participants of the Endoplasmic Reticulum-Associated Protein Degradation Machinery in the Human Mast Cell Leukemia Cell Line HMC-1.2. Cells 13, 747. 10.3390/cells13090747

Almanza, A., Carlesso, A., Chintha, C., Creedican, S., Doultsinos, D., Leuzzi, B., Luís, A., McCarthy, N., Montibeller, L., More, S., Papaioannou, A., Püschel, F., Sassano, M.L., Skoko, J., Agostinis, P., de Belleroche, J., Eriksson, L.A., Fulda, S., Gorman, A.M., Healy, S., Kozlov, A., Muñoz-Pinedo, C., Rehm, M., Chevet, E., Samali, A., 2019. Endoplasmic reticulum stress signalling - from basic mechanisms to clinical applications. FEBS J. 286, 241–278. 10.1111/febs.14608

Alonso-López, D., Campos-Laborie, F.J., Gutiérrez, M.A., Lambourne, L., Calderwood, M.A., Vidal, M., De Las Rivas, J., 2019. APID database: redefining protein–protein interaction experimental evidences and binary interactomes. Database J. Biol. Databases Curation 2019, baz005. 10.1093/database/baz005

Ashby, M.C., Tepikin, A.V., 2001. ER calcium and the functions of intracellular organelles. Semin. Cell Dev. Biol. 12, 11–17. 10.1006/scdb.2000.0212

Bartel, P., Chien, C.T., Sternglanz, R., Fields, S., 1993. Elimination of false positives that arise in using the two-hybrid system. BioTechniques 14, 920–924.

Bays, N.W., Gardner, R.G., Seelig, L.P., Joazeiro, C.A., Hampton, R.Y., 2001. Hrd1p/Der3p is a membrane-anchored ubiquitin ligase required for ER-associated degradation. Nat. Cell Biol. 3, 24–29. 10.1038/35050524

B’chir, W., Maurin, A.-C., Carraro, V., Averous, J., Jousse, C., Muranishi, Y., Parry, L., Stepien, G., Fafournoux, P., Bruhat, A., 2013. The eIF2α/ATF4 pathway is essential for stress-induced autophagy gene expression. Nucleic Acids Res. 41, 7683–7699. 10.1093/nar/gkt563

Belyy, V., Zuazo-Gaztelu, I., Alamban, A., Ashkenazi, A., Walter, P., 2022. Endoplasmic reticulum stress activates human IRE1α through reversible assembly of inactive dimers into small oligomers. eLife 11, e74342. 10.7554/eLife.74342

Binder, J.X., Pletscher-Frankild, S., Tsafou, K., Stolte, C., O’Donoghue, S.I., Schneider, R., Jensen, L.J., 2014. COMPARTMENTS: unification and visualization of protein subcellular localization evidence. Database 2014, bau012. 10.1093/database/bau012

Braakman, I., Hebert, D.N., 2013. Protein Folding in the Endoplasmic Reticulum. Cold Spring Harb. Perspect. Biol. 5, a013201–a013201. 10.1101/cshperspect.a013201

Cairrão, F., Santos, C.C., Le Thomas, A., Marsters, S., Ashkenazi, A., Domingos, P.M., 2022. Pumilio protects Xbp1 mRNA from regulated Ire1-dependent decay. Nat. Commun. 13, 1587. 10.1038/s41467-022-29105-x

Carreras-Sureda, A., Zhang, X., Laubry, L., Brunetti, J., Koenig, S., Wang, X., Castelbou, C., Hetz, C., Liu, Y., Frieden, M., Demaurex, N., 2023. The ER stress sensor IRE1 interacts with STIM1 to promote store-operated calcium entry, T cell activation, and muscular differentiation. Cell Rep. 42, 113540. 10.1016/j.celrep.2023.113540

Chen, R., Li, L., Weng, Z., 2003. ZDOCK: An initial-stage protein-docking algorithm. Proteins Struct. Funct. Bioinforma. 52, 80–87. 10.1002/prot.10389

Child, J.R., Chen, Q., Reid, D.W., Jagannathan, S., Nicchitta, C.V., 2021. Recruitment of endoplasmic reticulum-targeted and cytosolic mRNAs into membrane-associated stress granules. RNA 27, 1241–1256. 10.1261/rna.078858.121

Christoffer, C., Bharadwaj, V., Luu, R., Kihara, D., 2021. LZerD Protein-Protein Docking Webserver Enhanced With de novo Structure Prediction. Front. Mol. Biosci. 8. 10.3389/fmolb.2021.724947

Cloots, E., Guilbert, P., Provost, M., Neidhardt, L., Van de Velde, E., Fayazpour, F., De Sutter, D., Savvides, S.N., Eyckerman, S., Janssens, S., 2023. Activation of goblet-cell stress sensor IRE1β is controlled by the mucin chaperone AGR2. EMBO J. 1–24. 10.1038/s44318-023-00015-y

Colland, F., Jacq, X., Trouplin, V., Mougin, C., Groizeleau, C., Hamburger, A., Meil, A., Wojcik, J., Legrain, P., Gauthier, J.-M., 2004. Functional Proteomics Mapping of a Human Signaling Pathway. Genome Res. 14, 1324–1332. 10.1101/gr.2334104

Cox, J.S., Shamu, C.E., Walter, P., 1993. Transcriptional induction of genes encoding endoplasmic reticulum resident proteins requires a transmembrane protein kinase. Cell 73, 1197–1206. 10.1016/0092-8674(93)90648-a

Dufey, E., Bravo-San Pedro, J.M., Eggers, C., González-Quiroz, M., Urra, H., Sagredo, A.I., Sepulveda, D., Pihán, P., Carreras-Sureda, A., Hazari, Y., Sagredo, E.A., Gutierrez, D., Valls, C., Papaioannou, A., Acosta-Alvear, D., Campos, G., Domingos, P.M., Pedeux, R., Chevet, E., Alvarez, A., Godoy, P., Walter, P., Glavic, A., Kroemer, G., Hetz, C., 2020. Genotoxic stress triggers the activation of IRE1α-dependent RNA decay to modulate the DNA damage response. Nat. Commun. 11, 2401. 10.1038/s41467-020-15694-y

Durand, S., Franks, T.M., Lykke-Andersen, J., 2016. Hyperphosphorylation amplifies UPF1 activity to resolve stalls in nonsense-mediated mRNA decay. Nat. Commun. 7, 12434. 10.1038/ncomms12434

Fagone, P., Jackowski, S., 2009. Membrane phospholipid synthesis and endoplasmic reticulum function. J. Lipid Res. 50, S311–S316. 10.1194/jlr.R800049-JLR200

Fischer, J.W., Busa, V.F., Shao, Y., Leung, A.K.L., 2020. Structure-mediated RNA decay by UPF1 and G3BP1. Mol. Cell 78, 70–84.e6. 10.1016/j.molcel.2020.01.021

Fromont-Racine, M., Rain, J.-C., Legrain, P., 1997. Toward a functional analysis of the yeast genome through exhaustive two-hybrid screens. Nat. Genet. 16, 277–282. 10.1038/ng0797-277

Ghaemmaghami, S., Huh, W.-K., Bower, K., Howson, R.W., Belle, A., Dephoure, N., O’Shea, E.K., Weissman, J.S., 2003. Global analysis of protein expression in yeast. Nature 425, 737–741. 10.1038/nature02046

Gómez-Puerta, S., Ferrero, R., Hochstoeger, T., Zubiri, I., Chao, J., Aragón, T., Voigt, F., 2022. Live imaging of the co-translational recruitment of XBP1 mRNA to the ER and its processing by diffuse, non-polarized IRE1α. eLife 11, e75580. 10.7554/eLife.75580

Gu, F., Nguyên, D.T., Stuible, M., Dubé, N., Tremblay, M.L., Chevet, E., 2004. Protein-tyrosine phosphatase 1B potentiates IRE1 signaling during endoplasmic reticulum stress. J. Biol. Chem. 279, 49689–49693. 10.1074/jbc.C400261200

Guil, S., Long, J.C., Cáceres, J.F., 2006. hnRNP A1 Relocalization to the Stress Granules Reflects a Role in the Stress Response. Mol. Cell. Biol. 26, 5744–5758. 10.1128/MCB.00224-06

Guo, J., Guo, S., Lu, S., Gong, J., Wang, L., Ding, L., Chen, Q., Liu, W., 2023. The development of proximity labeling technology and its applications in mammals, plants, and microorganisms. Cell Commun. Signal. 21, 1–22. 10.1186/s12964-023-01310-1

Hanson, S.R., Culyba, E.K., Hsu, T.-L., Wong, C.-H., Kelly, J.W., Powers, E.T., 2009. The core trisaccharide of an N-linked glycoprotein intrinsically accelerates folding and enhances stability. Proc. Natl. Acad. Sci. 106, 3131–3136. 10.1073/pnas.0810318105

Harding, H.P., Zhang, Y., Zeng, H., Novoa, I., Lu, P.D., Calfon, M., Sadri, N., Yun, C., Popko, B., Paules, R., Stojdl, D.F., Bell, J.C., Hettmann, T., Leiden, J.M., Ron, D., 2003. An integrated stress response regulates amino acid metabolism and resistance to oxidative stress. Mol. Cell 11, 619–633. 10.1016/s1097-2765(03)00105-9

Haze, K., Yoshida, H., Yanagi, H., Yura, T., Mori, K., 1999. Mammalian transcription factor ATF6 is synthesized as a transmembrane protein and activated by proteolysis in response to endoplasmic reticulum stress. Mol. Biol. Cell 10, 3787–3799. 10.1091/mbc.10.11.3787

Heo, L., Lee, H., Seok, C., 2016. GalaxyRefineComplex: Refinement of protein-protein complex model structures driven by interface repacking. Sci. Rep. 6, 32153. 10.1038/srep32153

Hetz, C., Glimcher, L.H., 2009. Fine-Tuning of the Unfolded Protein Response: Assembling the IRE1α Interactome. Mol. Cell 35, 551–561. 10.1016/j.molcel.2009.08.021

Hetz, C., Zhang, K., Kaufman, R.J., 2020. Mechanisms, regulation and functions of the unfolded protein response. Nat. Rev. Mol. Cell Biol. 21, 421–438. 10.1038/s41580-020-0250-z

Hollien, J., Lin, J.H., Li, H., Stevens, N., Walter, P., Weissman, J.S., 2009. Regulated Ire1-dependent decay of messenger RNAs in mammalian cells. J. Cell Biol. 186, 323–331. 10.1083/jcb.200903014

Hollien, J., Weissman, J.S., 2006. Decay of Endoplasmic Reticulum-Localized mRNAs During the Unfolded Protein Response. Science 313, 104–107. 10.1126/science.1129631

Hughes, D., Mallucci, G.R., 2019. The unfolded protein response in neurodegenerative disorders – therapeutic modulation of the PERK pathway. FEBS J. 286, 342–355. 10.1111/febs.14422

Igbaria, A., Merksamer, P.I., Trusina, A., Tilahun, F., Johnson, J.R., Brandman, O., Krogan, N.J., Weissman, J.S., Papa, F.R., 2019. Chaperone-mediated reflux of secretory proteins to the cytosol during endoplasmic reticulum stress. Proc. Natl. Acad. Sci. 116, 11291–11298. 10.1073/pnas.1904516116

J⊘nson, L., Vikesaa, J., Krogh, A., Nielsen, L.K., Hansen, T. vO, Borup, R., Johnsen, A.H., Christiansen, J., Nielsen, F.C., 2007. Molecular Composition of IMP1 Ribonucleoprotein Granules *. Mol. Cell. Proteomics 6, 798–811. 10.1074/mcp.M600346-MCP200

Jiménez-García, B., Pons, C., Fernández-Recio, J., 2013. pyDockWEB: a web server for rigid-body protein–protein docking using electrostatics and desolvation scoring. Bioinformatics 29, 1698–1699. 10.1093/bioinformatics/btt262

Jurkin, J., Henkel, T., Nielsen, A.F., Minnich, M., Popow, J., Kaufmann, T., Heindl, K., Hoffmann, T., Busslinger, M., Martinez, J., 2014. The mammalian tRNA ligase complex mediates splicing of *XBP1* mRNA and controls antibody secretion in plasma cells. EMBO J. 33, 2922–2936. 10.15252/embj.201490332

Kedersha, N., Panas, M.D., Achorn, C.A., Lyons, S., Tisdale, S., Hickman, T., Thomas, M., Lieberman, J., McInerney, G.M., Ivanov, P., Anderson, P., 2016. G3BP–Caprin1–USP10 complexes mediate stress granule condensation and associate with 40S subunits. J. Cell Biol. 212, e201508028. 10.1083/jcb.201508028

Kelley, L.A., Gardner, S.P., Sutcliffe, M.J., 1996. An automated approach for clustering an ensemble of NMR-derived protein structures into conformationally related subfamilies. Protein Eng. 9, 1063–1065. 10.1093/protein/9.11.1063

Ko, J., Park, H., Heo, L., Seok, C., 2012. GalaxyWEB server for protein structure prediction and refinement. Nucleic Acids Res. 40, W294–W297. 10.1093/nar/gks493

Kozakov, D., Hall, D.R., Xia, B., Porter, K.A., Padhorny, D., Yueh, C., Beglov, D., Vajda, S., 2017. The ClusPro web server for protein–protein docking. Nat. Protoc. 12, 255–278. 10.1038/nprot.2016.169

Kroeger, H., Grimsey, N., Paxman, R., Chiang, W.-C., Plate, L., Jones, Y., Shaw, P.X., Trejo, J., Tsang, S.H., Powers, E., Kelly, J.W., Wiseman, R.L., Lin, J.H., 2018. The unfolded protein response regulator ATF6 promotes mesodermal differentiation. Sci. Signal. 11, eaan5785. 10.1126/scisignal.aan5785

Lajoie, P., Snapp, E.L., 2020. Size-dependent secretory protein reflux into the cytosol in association with acute endoplasmic reticulum stress. Traffic 21, 419–429. 10.1111/tra.12729

Le Goupil, S., Laprade, H., Aubry, M., Chevet, E., 2024. Exploring the IRE1 interactome: from canonical signaling functions to unexpected roles. J. Biol. Chem. 0. 10.1016/j.jbc.2024.107169

Lee, A.-H., Iwakoshi, N.N., Glimcher, L.H., 2003. XBP-1 Regulates a Subset of Endoplasmic Reticulum Resident Chaperone Genes in the Unfolded Protein Response. Mol. Cell. Biol. 23, 7448–7459. 10.1128/MCB.23.21.7448-7459.2003

Lee, A.-H., Scapa, E.F., Cohen, D.E., Glimcher, L.H., 2008. Regulation of hepatic lipogenesis by the transcription factor XBP1. Science 320, 1492–1496. 10.1126/science.1158042

Lhomond, S., Avril, T., Dejeans, N., Voutetakis, K., Doultsinos, D., McMahon, M., Pineau, R., Obacz, J., Papadodima, O., Jouan, F., Bourien, H., Logotheti, M., Jégou, G., Pallares-Lupon, N., Schmit, K., Le Reste, P., Etcheverry, A., Mosser, J., Barroso, K., Vauléon, E., Maurel, M., Samali, A., Patterson, J.B., Pluquet, O., Hetz, C., Quillien, V., Chatziioannou, A., Chevet, E., 2018. Dual IRE 1 RN ase functions dictate glioblastoma development. EMBO Mol. Med. 10. 10.15252/emmm.201707929

Liang, J., Yin, C., Doong, H., Fang, S., Peterhoff, C., Nixon, R.A., Monteiro, M.J., 2006. Characterization of erasin (UBXD2): a new ER protein that promotes ER-associated protein degradation. J. Cell Sci. 119, 4011–4024. 10.1242/jcs.03163

Liang, S., Tran, E., Du, X., Dong, J., Sudholz, H., Chen, H., Qu, Z., Huntington, N.D., Babon, J.J., Kershaw, N.J., Zhang, Z.-Y., Baell, J.B., Wiede, F., Tiganis, T., 2023. A small molecule inhibitor of PTP1B and PTPN2 enhances T cell anti-tumor immunity. Nat. Commun. 14, 4524. 10.1038/s41467-023-40170-8

Liu, S., Zhang, X., Yao, X., Wang, G., Huang, S., Chen, P., Tang, M., Cai, J., Wu, Z., Zhang, Y., Xu, R., Liu, K., He, K., Wang, Y., Jiang, L., Wang, Q.A., Rui, L., Liu, J., Liu, Y., 2024. Mammalian IRE1α dynamically and functionally coalesces with stress granules. Nat. Cell Biol. 1–15. 10.1038/s41556-024-01418-7

Longman, D., Jackson-Jones, K.A., Maslon, M.M., Murphy, L.C., Young, R.S., Stoddart, J.J., Hug, N., Taylor, M.S., Papadopoulos, D.K., Cáceres, J.F., 2020. Identification of a localized nonsense-mediated decay pathway at the endoplasmic reticulum. Genes Dev. 34, 1075–1088. 10.1101/gad.338061.120

Lu, C., Wu, C., Ghoreishi, D., Chen, W., Wang, L., Damm, W., Ross, G.A., Dahlgren, M.K., Russell, E., Von Bargen, C.D., Abel, R., Friesner, R.A., Harder, E.D., 2021. OPLS4: Improving Force Field Accuracy on Challenging Regimes of Chemical Space. J. Chem. Theory Comput. 17, 4291–4300. 10.1021/acs.jctc.1c00302

Madden, E., Logue, S.E., Healy, S.J., Manie, S., Samali, A., 2019. The role of the unfolded protein response in cancer progression: From oncogenesis to chemoresistance. Biol. Cell 111, 1–17. 10.1111/boc.201800050

Mahdizadeh, S.J., Thomas, M., Eriksson, L.A., 2021. Reconstruction of the Fas-Based Death-Inducing Signaling Complex (DISC) Using a Protein-Protein Docking Meta-Approach. J. Chem. Inf. Model. 61, 3543–3558. 10.1021/acs.jcim.1c00301

Marciniak, S.J., Yun, C.Y., Oyadomari, S., Novoa, I., Zhang, Y., Jungreis, R., Nagata, K., Harding, H.P., Ron, D., 2004. CHOP induces death by promoting protein synthesis and oxidation in the stressed endoplasmic reticulum. Genes Dev. 18, 3066–3077. 10.1101/gad.1250704

McGrath, E.P., Centonze, F.G., Chevet, E., Avril, T., Lafont, E., 2021. Death sentence: The tale of a fallen endoplasmic reticulum. Biochim. Biophys. Acta BBA - Mol. Cell Res. 1868, 119001. 10.1016/j.bbamcr.2021.119001

Mori, K., Ma, W., Gething, M.J., Sambrook, J., 1993. A transmembrane protein with a cdc2+/CDC28-related kinase activity is required for signaling from the ER to the nucleus. Cell 74, 743–756. 10.1016/0092-8674(93)90521-q

Novoa, I., Zeng, H., Harding, H.P., Ron, D., 2001. Feedback Inhibition of the Unfolded Protein Response by GADD34-Mediated Dephosphorylation of eIF2α. J. Cell Biol. 153, 1011–1022. 10.1083/jcb.153.5.1011

Offterdinger, M., Bastiaens, P.I., 2008. Prolonged EGFR Signaling by ERBB2-Mediated Sequestration at the Plasma Membrane. Traffic 9, 147–155. 10.1111/j.1600-0854.2007.00665.x

Olzmann, J.A., Kopito, R.R., Christianson, J.C., 2013. The Mammalian Endoplasmic Reticulum-Associated Degradation System. Cold Spring Harb. Perspect. Biol. 5, a013185–a013185. 10.1101/cshperspect.a013185

Orchard, S., Ammari, M., Aranda, B., Breuza, L., Briganti, L., Broackes-Carter, F., Campbell, N.H., Chavali, G., Chen, C., del-Toro, N., Duesbury, M., Dumousseau, M., Galeota, E., Hinz, U., Iannuccelli, M., Jagannathan, S., Jimenez, R., Khadake, J., Lagreid, A., Licata, L., Lovering, R.C., Meldal, B., Melidoni, A.N., Milagros, M., Peluso, D., Perfetto, L., Porras, P., Raghunath, A., Ricard-Blum, S., Roechert, B., Stutz, A., Tognolli, M., van Roey, K., Cesareni, G., Hermjakob, H., 2014. The MIntAct project—IntAct as a common curation platform for 11 molecular interaction databases. Nucleic Acids Res. 42, D358–D363. 10.1093/nar/gkt1115

Oughtred, R., Rust, J., Chang, C., Breitkreutz, B.-J., Stark, C., Willems, A., Boucher, L., Leung, G., Kolas, N., Zhang, F., Dolma, S., Coulombe-Huntington, J., Chatr-Aryamontri, A., Dolinski, K., Tyers, M., 2021. The BioGRID database: A comprehensive biomedical resource of curated protein, genetic, and chemical interactions. Protein Sci. Publ. Protein Soc. 30, 187–200. 10.1002/pro.3978

Papaioannou, A., Centonze, F., Metais, A., Maurel, M., Negroni, L., Gonzalez-Quiroz, M., Mahdizadeh, S.J., Svensson, G., Zare, E., Blondel, A., Koong, A.C., Hetz, C., Pedeux, R., Tremblay, M.L., Eriksson, L.A., Chevet, E., 2022. Stress-induced tyrosine phosphorylation of RtcB modulates IRE1 activity and signaling outputs. Life Sci. Alliance 5, e202201379. 10.26508/lsa.202201379

Rastelli, G., Rio, A.D., Degliesposti, G., Sgobba, M., 2010. Fast and accurate predictions of binding free energies using MM-PBSA and MM-GBSA. J. Comput. Chem. 31, 797–810. 10.1002/jcc.21372

Sassano, M.L., Derua, R., Waelkens, E., Agostinis, P., van Vliet, A.R., 2021. Interactome Analysis of the ER Stress Sensor Perk Uncovers Key Components of ER-Mitochondria Contact Sites and Ca2+ Signalling. Contact 4, 25152564211052392. 10.1177/25152564211052392

Schröder, M., Kaufman, R.J., 2005. THE MAMMALIAN UNFOLDED PROTEIN RESPONSE. Annu. Rev. Biochem. 74, 739–789. 10.1146/annurev.biochem.73.011303.074134

Sears, R.M., May, D.G., Roux, K.J., 2019. BioID as a Tool for Protein-Proximity Labeling in Living Cells. Methods Mol. Biol. Clifton NJ 2012, 299–313. 10.1007/978-1-4939-9546-2_15

Shannon, P., Markiel, A., Ozier, O., Baliga, N.S., Wang, J.T., Ramage, D., Amin, N., Schwikowski, B., Ideker, T., 2003. Cytoscape: a software environment for integrated models of biomolecular interaction networks. Genome Res. 13, 2498–2504. 10.1101/gr.1239303

Shoulders, M.D., Ryno, L.M., Genereux, J.C., Moresco, J.J., Tu, P.G., Wu, C., Yates, J.R., Su, A.I., Kelly, J.W., Wiseman, R.L., 2013. Stress-Independent Activation of XBP1s and/or ATF6 Reveals Three Functionally Diverse ER Proteostasis Environments. Cell Rep. 3, 1279–1292. 10.1016/j.celrep.2013.03.024

Sicari, D., Centonze, F.G., Pineau, R., Le Reste, P.-J., Negroni, L., Chat, S., Mohtar, M.A., Thomas, D., Gillet, R., Hupp, T., Chevet, E., Igbaria, A., 2021. Reflux of Endoplasmic Reticulum proteins to the cytosol inactivates tumor suppressors. EMBO Rep. 22, e51412. 10.15252/embr.202051412

Sidrauski, C., Walter, P., 1997. The Transmembrane Kinase Ire1p Is a Site-Specific Endonuclease That Initiates mRNA Splicing in the Unfolded Protein Response. Cell 90, 1031–1039. 10.1016/S0092-8674(00)80369-4

Sun, S., Shi, G., Sha, H., Ji, Y., Han, X., Shu, X., Ma, H., Inoue, T., Gao, B., Kim, H., Bu, P., Guber, R.D., Shen, X., Lee, A.-H., Iwawaki, T., Paton, A.W., Paton, J.C., Fang, D., Tsai, B., Yates III, J.R., Wu, H., Kersten, S., Long, Q., Duhamel, G.E., Simpson, K.W., Qi, L., 2015. IRE1α is an endogenous substrate of endoplasmic-reticulum-associated degradation. Nat. Cell Biol. 17, 1546–1555. 10.1038/ncb3266

Sundaram, A., Plumb, R., Appathurai, S., Mariappan, M., 2017. The Sec61 translocon limits IRE1α signaling during the unfolded protein response. eLife 6, e27187. 10.7554/eLife.27187

Szklarczyk, D., Gable, A.L., Lyon, D., Junge, A., Wyder, S., Huerta-Cepas, J., Simonovic, M., Doncheva, N.T., Morris, J.H., Bork, P., Jensen, L.J., Mering, C. von, 2019. STRING v11: protein-protein association networks with increased coverage, supporting functional discovery in genome-wide experimental datasets. Nucleic Acids Res. 47, D607–D613. 10.1093/nar/gky1131

Thomas, P.D., Ebert, D., Muruganujan, A., Mushayahama, T., Albou, L.-P., Mi, H., 2022. PANTHER: Making genome-scale phylogenetics accessible to all. Protein Sci. 31, 8–22. 10.1002/pro.4218

Tong, L., 2013. Structure and function of biotin-dependent carboxylases. Cell. Mol. Life Sci. CMLS 70, 863–891. 10.1007/s00018-012-1096-0

Tovchigrechko, Andrey, Vasker, Ilya, 2006. GRAMM-X public web server for protein-protein docking - PubMed [WWW Document]. URL https://pubmed.ncbi.nlm.nih.gov/16845016/ (accessed 4.21.24).

Tyanova, S., Temu, T., Sinitcyn, P., Carlson, A., Hein, M.Y., Geiger, T., Mann, M., Cox, J., 2016. The Perseus computational platform for comprehensive analysis of (prote)omics data. Nat. Methods 13, 731–740. 10.1038/nmeth.3901

Uezu, A., Kanak, D.J., Bradshaw, T.W.A., Soderblom, E.J., Catavero, C.M., Burette, A.C., Weinberg, R.J., Soderling, S.H., 2016. Identification of an elaborate complex mediating postsynaptic inhibition. Science 353, 1123–1129. 10.1126/science.aag0821

Upton, J.-P., Wang, L., Han, D., Wang, E.S., Huskey, N.E., Lim, L., Truitt, M., McManus, M.T., Ruggero, D., Goga, A., Papa, F.R., Oakes, S.A., 2012. IRE1α Cleaves Select microRNAs During ER Stress to Derepress Translation of Proapoptotic Caspase-2. Science 338, 818–822. 10.1126/science.1226191

Urra, H., Henriquez, D.R., Cánovas, J., Villarroel-Campos, D., Carreras-Sureda, A., Pulgar, E., Molina, E., Hazari, Y.M., Limia, C.M., Alvarez-Rojas, S., Figueroa, R., Vidal, R.L., Rodriguez, D.A., Rivera, C.A., Court, F.A., Couve, A., Qi, L., Chevet, E., Akai, R., Iwawaki, T., Concha, M.L., Glavic, Á., Gonzalez-Billault, C., Hetz, C., 2018. IRE1α governs cytoskeleton remodelling and cell migration through a direct interaction with filamin A. Nat. Cell Biol. 20, 942–953. 10.1038/s41556-018-0141-0

van Vliet, A.R., Giordano, F., Gerlo, S., Segura, I., Van Eygen, S., Molenberghs, G., Rocha, S., Houcine, A., Derua, R., Verfaillie, T., Vangindertael, J., De Keersmaecker, H., Waelkens, E., Tavernier, J., Hofkens, J., Annaert, W., Carmeliet, P., Samali, A., Mizuno, H., Agostinis, P., 2017. The ER Stress Sensor PERK Coordinates ER-Plasma Membrane Contact Site Formation through Interaction with Filamin-A and F-Actin Remodeling. Mol. Cell 65, 885–899.e6. 10.1016/j.molcel.2017.01.020

Vattem, K.M., Wek, R.C., 2004. Reinitiation involving upstream ORFs regulates ATF4 mRNA translation in mammalian cells. Proc. Natl. Acad. Sci. 101, 11269–11274. 10.1073/pnas.0400541101

Wall, M.L., Bera, A., Wong, F.K., Lewis, S.M., 2020. Cellular stress orchestrates the localization of hnRNP H to stress granules. Exp. Cell Res. 394, 112111. 10.1016/j.yexcr.2020.112111

Welihinda, A.A., Tirasophon, W., Green, S.R., Kaufman, R.J., 1998. Protein Serine/Threonine Phosphatase Ptc2p Negatively Regulates the Unfolded-Protein Response by Dephosphorylating Ire1p Kinase. Mol. Cell. Biol. 18, 1967–1977. 10.1128/MCB.18.4.1967

Wu, J., Rutkowski, D.T., Dubois, M., Swathirajan, J., Saunders, T., Wang, J., Song, B., Yau, G.D.-Y., Kaufman, R.J., 2007. ATF6α Optimizes Long-Term Endoplasmic Reticulum Function to Protect Cells from Chronic Stress. Dev. Cell 13, 351–364. 10.1016/j.devcel.2007.07.005

Yamamoto, K., Suzuki, N., Wada, T., Okada, T., Yoshida, H., Kaufman, R.J., Mori, K., 2008. Human HRD1 Promoter Carries a Functional Unfolded Protein Response Element to Which XBP1 but not ATF6 Directly Binds. J. Biochem. (Tokyo) 144, 477–486. 10.1093/jb/mvn091

Yan, Y., Tao, H., He, J., Huang, S.-Y., 2020. The HDOCK server for integrated protein– protein docking. Nat. Protoc. 15, 1829–1852. 10.1038/s41596-020-0312-x

Yang, Z., Zhang, J., Jiang, D., Khatri, P., Solow-Cordero, D.E., Toesca, D.A.S., Koumenis, C., Denko, N.C., Giaccia, A.J., Le, Q.-T., Koong, A.C., 2018. A Human Genome-Wide RNAi Screen Reveals Diverse Modulators that Mediate IRE1α– XBP1 Activation. Mol. Cancer Res. 16, 745–753. 10.1158/1541-7786.MCR-17-0307

Yildirim, A.D., Citir, M., Dogan, A.E., Veli, Z., Yildirim, Z., Tufanli, O., Traynor-Kaplan, A., Schultz, C., Erbay, E., 2022. ER Stress-Induced Sphingosine-1-Phosphate Lyase Phosphorylation Potentiates the Mitochondrial Unfolded Protein Response. J. Lipid Res. 63, 100279. 10.1016/j.jlr.2022.100279

Yildirim, Z., Baboo, S., Hamid, S.M., Dogan, A.E., Tufanli, O., Robichaud, S., Emerton, C., Diedrich, J.K., Vatandaslar, H., Nikolos, F., Gu, Y., Iwawaki, T., Tarling, E., Ouimet, M., Nelson, D.L., Yates, J.R., Walter, P., Erbay, E., 2022. Intercepting IRE1 kinase-FMRP signaling prevents atherosclerosis progression. EMBO Mol. Med. 14. 10.15252/emmm.202115344

