## Supplementary material for "Proximity Interactome analyses unveil novel regulators of IRE1α canonical signaling": LeGoupil_et_al_Suppl

**Table S1:** List of siRNAs used for knock-down experiments.

| siRNA | Reference |
| --- | --- |
| **siCtrl** | D-001210-02-05 |
| **HNRNPL** | M-011293-01-0005 |
| **RPN1** | M-018903-01-0005 |
| **PDIA4** | M-019249-01-0005 |
| **PTPN1** | M-003529-04-0005 |
| **PHB** | M-010530-00-0005 |
| **UPF1** | M-011763-01-0005 |
| **COPB1** | M-017940-00-0005 |

**Table S2:** List of antibodies used for western blot.

| Antibody | Company/Reference | Working dilution |
|---|---|---|
| **Anti-Mouse HRP** | CST/7076 | 1:7000 |
| **Anti-Rabbit HRP** | CST/7074 | 1:7000 |
| **Calnexin** | Gift from JJM Bergeron (McGill, Canada) (Ou *et al*, 1993) | 1:2000 |
| **COPB1** | SantaCruz/sc-165976 | 1:2000 |
| **Flag** | CST/2368 | 1:2000 |
| **HNRNPL** | CST/65043 | 1:2000 |
| **IRE1α** | CST/3294 | 1:2000 |
| **PDIA4** | CST/2798 | 1:1000 |
| **PTPN1** | SantaCruz/sc-56960 | 1:2000 |
| **Streptavidin-HRP** | Abcam/ab7403 | 1:3000 |
| **SYVN1** | CST14773 | 1:2000 |
| **XBP1s** | ProteinTech/24868 | 1:1000 |
| **β-actin** | CST/3700 | 1:4000 |

**Table S3:** List of primers used for RT-qPCR.

| Name | Forward primer (5'-3') | Reverse primer (5'-3') |
|---|---|---|
| **CD59** | GCC TGC ACT GCT ACA ACT | CAA TGC TCA AAC TTC CAA CAC T |
| **CHOP** | ATT GAC CGA ATG GTG AAT CTG C | AGC TGA GAC CTT TCC TTT TGT CTA |
| **COPB1** | AAC TCC TAG TGC GAA CAT TGC | CTG CTG CTT CGT TGT TGT CAC |
| **ERDJ4** | GCCATGAAGTACCACCCTGACA | TCGCTATTAGCATCTGAGAGTGT |
| **HERPUD1** | AAC GGC ATG TTT TGC ATC TG | GGG GAA GAA AGG TTC CGA AG |
| **HNRNPL** | TAC GCA GCC GAC AAC CAA ATA | CTC CGG GAG TCA TCC GAG T |
| **IRE1** | AGA GAA GCA GCA GAC TTT GTC | GTT TTG GTG TCG TAC ATG GTG A |
| **PDIA4** | GGC AGG CTG TAG ACT ACG AG | TTG GTC AAC ACA AGC GTG ACT |
| **PHB** | GAC CAC GTA ATG TGC CAG TCA | CAT CAT AGT CCT CTC CGA TGC T |
| **PTPN1** | TCC CTT TGA CCA TAG TCG GAT | GTG ACC GCA TGT GTT AGG CA |
| **RPN1** | GGC CAA GAT TTC AGT CAT TGT GG | CTT CGT TGG ATA GGG AGA GTA GA |
| **SEL1L** | CAA TGC TTC CTT GTG CCG | AGG ACC CTT GGA TCA GTG GTC |
| **UPF1** | CTG CAA CGG ACG TGG AAA TAC | ACA GCC GCA GTT GTA GCA C |
| **XBP1s** | TGC TGA GTC CGC AGC AGG TG | GCT GGC AGG CTC TGG GGA AG |

#### Supplementary Figure Legends

**Figure S1: Validation of IRE1 KO in A375-MA2 cell line.** Western-Blot of IRE1 protein expression in A375-MA2 WT or KO for IRE1. One experiment representative of XX is shown. One representative experiment out of three is shown.

**Figure S2: Computational analysis of IRE1 interactome. A)** Venn diagram crossing the list of interactors found upon the two different ER stressor treatment Tunicamycin or Thapsigargin. **B)** Bar graph representing the proportion of basal, stress or unaltered interactors within each biological function. **C)** Bar graph representing the proportion of interactors within several subcellular compartments for the two interactome strategies (reference and BioID). Subcellular localization of a protein was assessed by the COMPARTMENTS database with a cut-off set at 4,75 (95%). **D)** Venn diagram crossing the list of interactors from reference and BioID IRE1 interactomes (left). Gene Ontology enrichment (Biological process) performed on the 49 common interactors (right). **E)** Venn diagram crossing the list of interactors from IRE1 and PERK BioIDs (left). Gene Ontology enrichment (Biological process) performed on the 43 common interactors (right). **F)** Venn diagram crossing the list of interactors from IRE1α and IRE1β interactomes. The 10 common interactors are listed on the right.

**Figure S3: IRE1 signalosome and direct interaction analysis. A)** Venn diagram crossing the list of interactors from reference and BioID IRE1 signalosomes. The 12 common signalosome partners are listed on the right. **B)** Table comparing the enrichment of signalosome proteins between the two interactome approaches (reference or BioID). A $\chi^2$ enrichment test was performed to evaluate the differential enrichment. **C)** Table representing the direct interaction prediction analysis between the proteins of interest. **D)** Summary table of the yeast-two-hybrid screening of direct interaction between IRE1 and the selected proteins of interest.

**Figure S4: Effect of proteins of interest downregulation on ER stress parameters. A)** RT-qPCR analysis of HNRNPL, RPN1, PDIA4, PTPN1, PHB, UPF1, COPB1 genes expression in HEK293T cells transfected with 50 nM siRNAs targeting each of the genes of interest for 48 hours. **B, C, D, E)** RT-qPCR analysis of ERDJ4, CHOP, SEL1L and HERPUD1 gene expression respectively in HEK293T cells transfected with 50 nM siRNAs targeting each of the genes of interest for 48 hours. **F)** Schematic representation of the UPR markers impacted by proteins of interest downregulation as evaluated in (B, C, D, E). (*P < 0.05, **P < 0.01, ***P < 0.001).

**Figure S5: Validation of HNRNPL-mediated expression of IRE1 in different cell lines.** Western Blot of IRE1 protein under basal conditions upon siCtrl or siHNRNPL-mediated silencing in 6 different cell lines (HEK293T, HeLa, MDA-MB-231, SUM 159, U87 and U251).

### Supplementary Figures

**Figure S1**

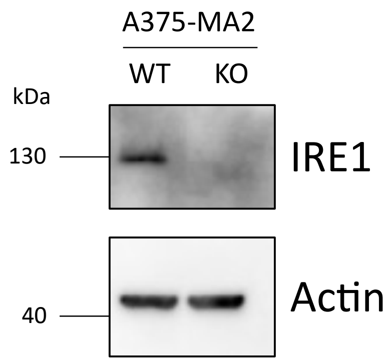

### Figure S2

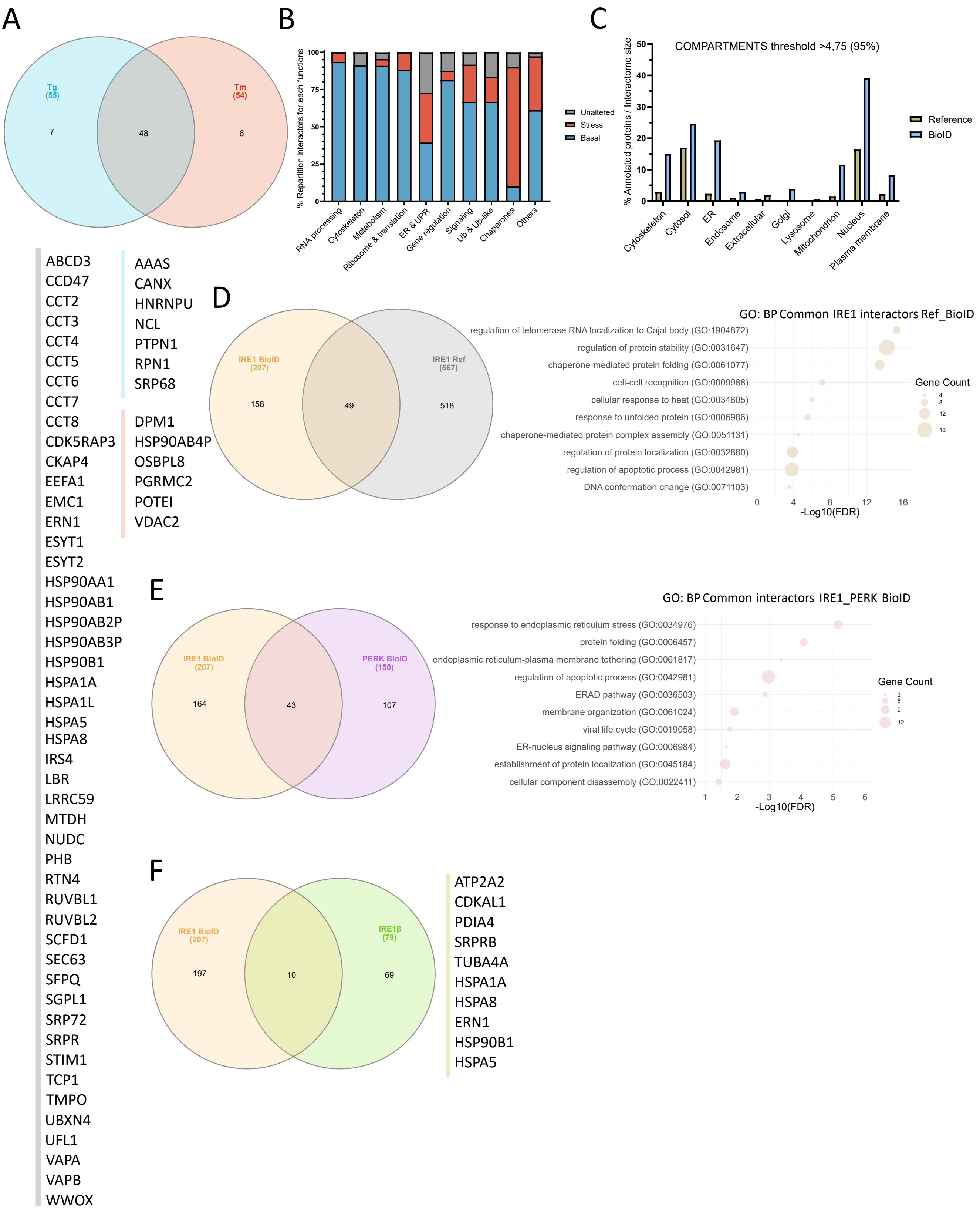
A
Tg
(55)
7
48
Tm
(54)
6
B
% Repartition interactors for each functions
100
75
50
25
0
RNA processing
Cytoskeleton
Metabolism
Ribosome & translation
ER & UPR
Gene regulation
Signaling
Ub & Ub-like
Chaperones
Others
Unaltered
Stress
Basal
C
COMPARTMENTS threshold >4,75 (95%)
% Annotated proteins / Interactome size
50
40
30
20
10
0
Cytoskeleton
Cytosol
ER
Endosome
Extracellular
Golgi
Lysosome
Mitochondrion
Nucleus
Plasma membrane
Reference
BioID
ABCD3
CCD47
CCT2
CCT3
CCT4
CCT5
CCT6
CCT7
CCT8
CDK5RAP3
CKAP4
EEFA1
EMC1
ERN1
ESYT1
ESYT2
HSP90AA1
HSP90AB1
HSP90AB2P
HSP90AB3P
HSP90B1
HSPA1A
HSPA1L
HSPA5
HSPA8
IRS4
LBR
LRRC59
MTDH
NUDC
PHB
RTN4
RUVBL1
RUVBL2
SCFD1
SEC63
SFPQ
SGPL1
SRP72
SRPR
STIM1
TCP1
TMPO
UBXN4
UFL1
VAPA
VAPB
WWOX
AAAS
CANX
HNRNPU
NCL
PTPN1
RPN1
SRP68
DPM1
HSP90AB4P
OSBPL8
PGRMC2
POTEI
VDAC2
D
IRE1 BioID
(207)
158
49
IRE1 Ref
(567)
518
GO: BP Common IRE1 interactors Ref_BioID
regulation of telomerase RNA localization to Cajal body (GO:1904872)
regulation of protein stability (GO:0031647)
chaperone-mediated protein folding (GO:0061077)
cell-cell recognition (GO:0009988)
cellular response to heat (GO:0034605)
response to unfolded protein (GO:0006986)
chaperone-mediated protein complex assembly (GO:0051131)
regulation of protein localization (GO:0032880)
regulation of apoptotic process (GO:0042981)
DNA conformation change (GO:0071103)
0
4
8
12
16
-Log10(FDR)
Gene Count
4
8
12
16
E
IRE1 BioID
(207)
164
43
PERK BioID
(150)
107
GO: BP Common interactors IRE1_PERK BioID
response to endoplasmic reticulum stress (GO:0034976)
protein folding (GO:0006457)
endoplasmic reticulum-plasma membrane tethering (GO:0061817)
regulation of apoptotic process (GO:0042981)
ERAD pathway (GO:0036503)
membrane organization (GO:0061024)
viral life cycle (GO:0019058)
ER-nucleus signaling pathway (GO:0006984)
establishment of protein localization (GO:0045184)
cellular component disassembly (GO:0022411)
1
2
3
4
5
6
-Log10(FDR)
Gene Count
3
6
9
12
F
IRE1 BioID
(207)
197
10
IRE1β
(79)
69
ATP2A2
CDKAL1
PDIA4
SRPRB
TUBA4A
HSPA1A
HSPA8
ERN1
HSP90B1
HSPA5

### Figure S3

A

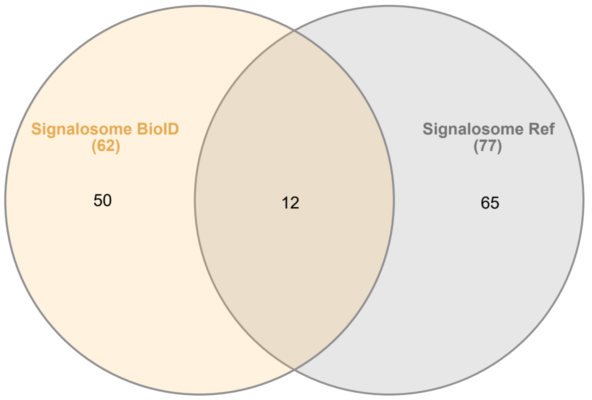


CCT4
CCT7
CHERP
ERN1
HSP90B1
HSPA5
PHB
RPS14
SEC63
SFPQ
UPF1
VC

B

| | Reference | BioID | $X^2$ test |
|---|---|---|---|
| Signalosome size | 77 | 62 | |
| Interactome size | 511 | 207 | p-value= $5.57x10^{-7}$ |
| % Signalosome /Interactome | 15% | 30% | |

C

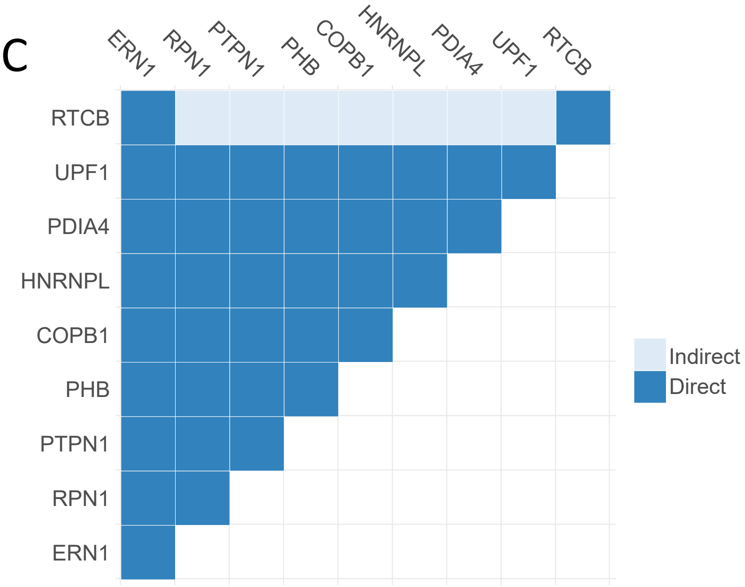


D

| Interaction Matrix | | Selection Medium | |
|---|---|---|---|
| Bait | Prey | DO-2 | DO-3 |
| ERN1 K599A | HNRNPL | + | - |
| | RPN1 | + | - |
| | PDIA4 | + | - |
| | PTPN1 | + | - |
| | UPF1 | + | - |
| | COPB1 | + | + |

### Figure S4

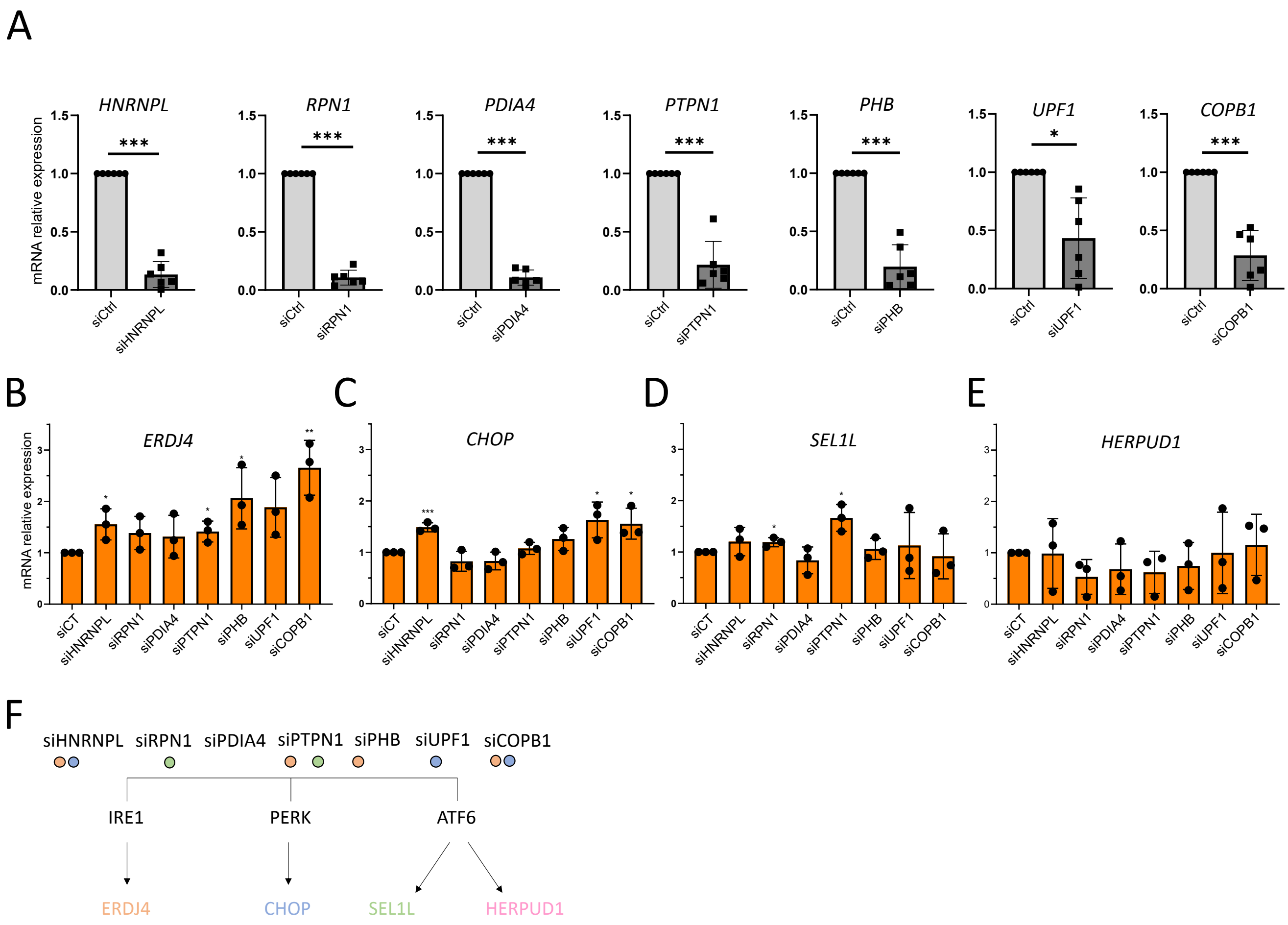
A
HNRNPL
RPN1
PDIA4
PTPN1
PHB
UPF1
COPB1
mRNA relative expression
siCtrl
siHNRNPL
siRPN1
siPDIA4
siPTPN1
siPHB
siUPF1
siCOPB1
B
ERDJ4
C
CHOP
D
SEL1L
E
HERPUD1
siCT
F
siHNRNPL siRPN1 siPDIA4 siPTPN1 siPHB siUPF1 siCOPB1
IRE1
PERK
ATF6
ERDJ4
CHOP
SEL1L
HERPUD1

### Figure S5

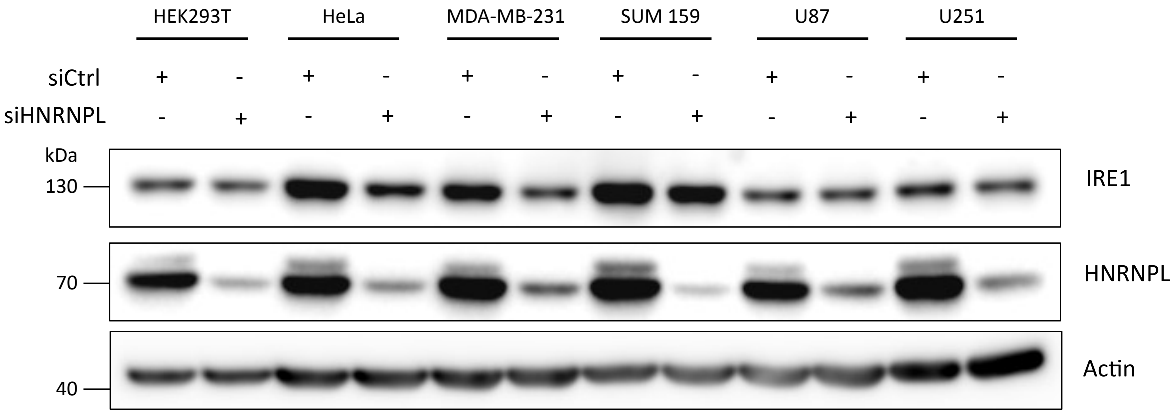
HEK293T
HeLa
MDA-MB-231
SUM 159
U87
U251
siCtrl
siHNRNPL
+
-
kDa
130
70
40
IRE1
HNRNPL
Actin
